# A Dimer-Seeded Pathway Governs the Nucleation of ProT*α*-H1 Biomolecular Condensates

**DOI:** 10.64898/2026.03.27.714715

**Authors:** Eduardo Pedraza, Andrés R. Tejedor, Álvaro S. Zorita, Rosana Collepardo-Guevara, David de Sancho, Pablo Llombart, Jorge R. Espinosa

## Abstract

Condensation between highly charged, intrinsically disordered proteins underlies physiologically essential processes such as chromatin regulation and nucleolus self-assembly, yet the mechanism by which these condensates first nucleate from the dilute phase has remained essentially inaccessible to both experiment and atomistic simulation. Here we use residue-level coarse-grained simulations of complex coacervates formed by Prothymosin *α* (ProT*α*) with histone H1, protamine, poly-lysine and poly-arginine, and show that they recapitulate the experimental salt-dependent phase behaviour, binding affinities, chain reconfiguration, protein diffusion and viscosity of these systems. Combining dilute-phase binding free energies with brute-force nucleation simulations, we reveal that these condensates assemble through a dimer-seeded nucleation mechanism: oppositely charged, polyanionic and polycationic partners first pair into a charge-compensated dimer—the most stable minimal complex and the dominant pre-nucleation species—from which clusters subsequently grow into large-scale condensates. The free energy cost of forming these clusters rises with salt and becomes density-controlled for larger assemblies, placing nucleation under the joint control of electrostatic screening and protein supersaturation. This same electrostatic screening shapes mature mesoscopic condensates: contacts form and break within picoseconds regardless of the phase, while chain reconfiguration and diffusion slow down markedly upon condensation, so that a condensate is set apart not by the strength of its contacts but by their collective network connectivity.

## I. INTRODUCTION

Cellular organization rests on the formation of membrane-bound and membrane-less organelles. The latter—e.g., biomolecular condensates—concentrate proteins and nucleic acids within the cytoplasm and nucleus, and control processes ranging from gene expression to stress response and signal transduction^1–7^. Representative examples include nucleoli, Cajal bodies, stress granules and P bodies. The ability of condensates to assemble and disassemble reversibly is tied to their material properties spanning fluid to viscoelastic regimes^8,9^. Disruption of this balance can produce aberrant, solid-like structures, a phenomenon implicated in several neurodegenerative disorders characterized by irreversible protein accumulation^10–13^. Linking condensate phase behaviour to its molecular origin therefore requires connecting the pathway of self-assembly—from initial dilute-phase encounters to the conformational rearrangement, diffusion and viscosity of the mature condensate—to the underlying network of intermolecular interactions. This network is, in turn, set by amino acid sequence, charge patterning and solution conditions such as temperature, concentration, pH and salt^14–25^.

Mixtures of oppositely charged intrinsically disordered proteins (IDPs) and nucleic acids phase-separate via complex coacervation and are a well-characterized model system for probing how nanoscale molecular dynamics give rise to mesoscale condensate formation and material properties^15,16,26–31^. In these systems, multivalent electrostatic interactions directly couple the molecular contact network to emergent rheological behaviour, and the intermolecular connectivity is highly tunable through salt concentration, which modulates both stability and viscosity^15–17,29^. Single-molecule Förster Resonance Energy Transfer (FRET) with nanosecond fluorescence correlation spectroscopy has resolved conformational dynamics and translational diffusion in these systems^32–34^, while microrheology, optical microscopy and particle tracking have quantified their viscoelastic properties and condensate dynamics^17,18,29,35–38^. However, directly linking residue-level interactions to emergent viscoelasticity, condensate stability and the nucleation process itself remains largely beyond the reach of most experimental approaches. In particular, the microscopic origin of condensate nucleation itself—*i*.*e*. how oppositely charged, highly disordered chains nucleate from the dilute phase into a mesoscopic dense phase—remains largely unexplored, even though this process sets the initial network topology and, ultimately, the material properties of mature condensates. In this respect, complex coacervates of oppositely charged IDPs closely parallel simple electrolyte solutions in water, in which association between oppositely charged species drives self-assembly and, beyond a critical supersaturation, the homogeneous nucleation of a new phase^39–42^. Condensate formation is an activated process: small clusters continuously form and redissolve in the dilute supersaturated phase, and only those that exceed a critical cluster size grow into a macroscopic dense phase^41,43^. Establishing this link between how a condensate first nucleates and the mesoscale properties it subsequently acquires—including its viscosity and the exchange of material between dense and dilute phases—would make possible to predict these properties directly from the molecular assembly pathway, rather than measuring them only once the condensate is already formed.

All-atom simulations with explicit solvent have been highly successful at resolving protein conformational reconfiguration, local diffusion, and intermolecular contact lifetimes in model complex coacervates such as in Prothymosin *α* (ProT*α*)–Histone 1 (H1) condensates^15,16,26,27^, providing molecular-level insight that is hardly accessible experimentally. However, their computational cost restricts accessible timescales, which precludes the observation of the initial stages of condensate formation itself: no mechanistic information on how condensation takes place is therefore accessible from all-atom simulations. Resolving the phase diagram across multiple salt and concentration conditions is similarly prohibitive, requiring elaborate multi-stage system preparation—typically an initial coarse-grained run, atomistic reconstruction, implicit-solvent pre-equilibration, and subsequent cloning and solvation^44–46^. Sequence-dependent coarse-grained (CG) models trade some molecular detail for access to full phase diagrams and larger spatiotemporal scales while retaining the sequence-specific interactions that govern phase behaviour and material properties such as viscosity, diffusion, surface tension and coarsening dynamics^47–54^.

Here we use the residue-level CG model Mpipi-Recharged^48,55,56^, an implicit-solvent, implicit-ion framework that improves on the original Mpipi model^57^ through a refined, pair-specific treatment of electrostatics and salt screening, to characterize the salt-dependent phase behaviour, dynamics and nucleation of ProT*α*-based complex coacervates. Condensate material properties, in this context, provide a stringent experimental benchmark for coarse-grained models, since they probe emergent, collective behaviour rather than the pairwise interactions the models are directly parametrized against. We first quantify, focusing on ProT*α* as the common component across systems, the phase diagram, conformational reconfiguration and translational diffusion that link sequence-dependent interactions to condensate viscosity^16^, together with residue-level monomeric mobility, and compare dynamics across dense and dilute phases to reveal the coexistence of fast intermolecular interactions with slow mesoscale relaxation at condensate-level, depending on molecular length scale. In parallel, having established the phase diagrams of these mixtures, we combine dilute-phase binding free energies for ProT*α*-H1 complexes^27^ with brute-force nucleation simulations^39^ to characterize how minimal complexes assemble into condensate-forming clusters. By performing spontaneous unseeded nucleation simulations of a supersaturated solution, we reveal a dimer-seeded nucleation pathway of ProT*α*-H1 condensates, linking the initial dimerization stage and the growth into a condensed phase. Compared against the phase diagrams, these simulations place nucleation on the same footing as the thermodynamics and dynamics of the condensed phase: they identify a charge-compensated ProT*α*-H1 dimer as the species from which nucleation proceeds, and show that the same electrostatic screening that controls the free-energy cost of forming larger clusters also sets the density, connectivity and viscoelasticity of the condensate that grows from them. This coupling between how a condensate forms and the material state it adopts over time is directly relevant to how condensation and dysregulation may occur *in vivo*.

## II. RESULTS

### A residue-resolution model near quantitively captures the experimental salt-dependent phase behaviour of complex coacervates

Complex coacervates form through association of oppositely charged polyelectrolytes, driven enthalpically by electrostatic pairing and entropically by counterion release and solvent reorganization. We investigate coacervates formed by negatively charged ProT*α* (net charge *−*43) with H1 (lysine-rich, +54), protamine (arginine-rich, +21), K50 and R50 (both +50) (Fig. 1a), all previously characterized *in vitro*^15,16^. Since complex coacervation is primarily electrostatically driven, stability is typically maximized close to stoichiometries where the total charges of the two components balance^16,58–60^. We therefore prepare mixtures at compositions close to electroneutrality (H1:ProT*α* 1:1.26, K50:ProT*α* 1:1.16, protamine:ProT*α* 1:0.49, R50:ProT*α* 1:1.16 molecule ratios; see Supplementary Section S2) as described in Ref.^16^. This design allows us to systematically probe how sequence composition, charge patterning and salt concentration shape condensate stability and dynamics, which we first use to validate the model before turning to the central question of how these condensates nucleate from solution.

**FIG. 1.**
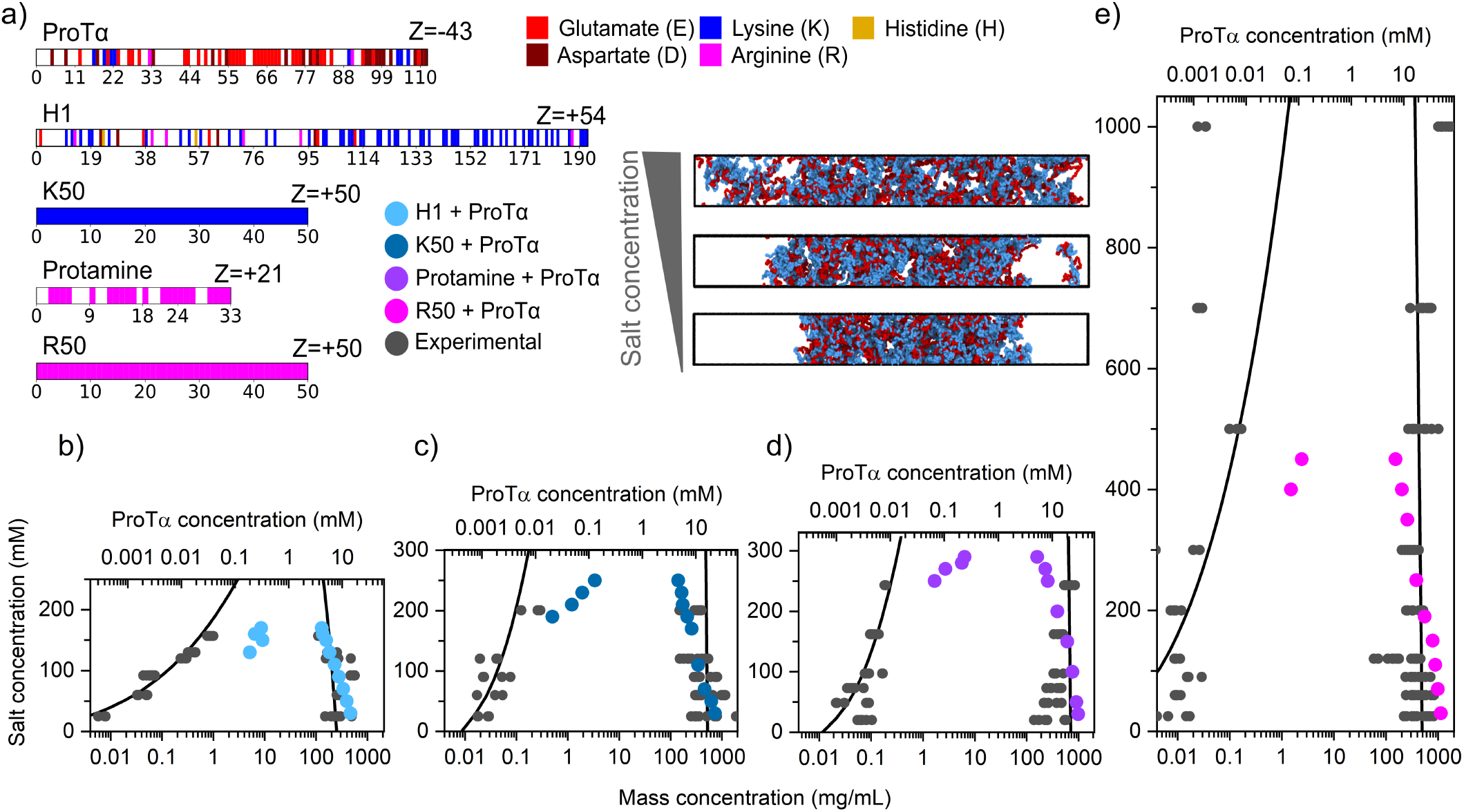
Mpipi-Recharged captures the salt-dependent phase behaviour of highly charged complex coacervates. (a) Charge distribution along the sequences of ProT*α*, H1, protamine, R50 and K50, with net charges *Z* (colour code: lysine blue, arginine magenta, glutamic acid red, aspartic acid maroon, histidine light orange), and representative Direct Coexistence simulation snapshots of ProT*α*-H1 at increasing monovalent salt. (b-e) Phase diagrams from Direct Coexistence simulations for H1+ProT*α* (1:1.26, light blue), K50+ProT*α* (1:1.16, dark blue), protamine+ProT*α* (1:0.49, violet) and R50+ProT*α* (1:1.16, magenta), with *in vitro* data (grey points) and the corresponding binodal fits (black lines)^15,16^. Total mass concentration is on the bottom axis, ProT*α* concentration on the top axis.

Using Direct Coexistence (DC) simulations^61–64^ with the Mpipi-Recharged residue-resolution force field^48^ (Supplementary Section S1), in which the dense and dilute phases coexist explicitly across an interface within an elongated simulation box, we compute the phase diagrams for these mixtures at *T* = 295 K as a function of monovalent salt concentration (Fig. 1a-e; Supplementary Section S3). Representative DC snapshots of ProT*α*-H1 at increasing salt concentration are shown in Fig. 1a. Our CG simulations near quantitatively reproduce the experimental dense-phase coexistence concentrations and capture the relative stability ordering of ProT*α*-H1, ProT*α*-K50, ProT*α*-protamine and ProT*α*-R50, as well as the experimental salt range over which ProT*α*-H1, ProT*α*-K50 and ProT*α*-protamine phase-separate. Dilute-phase saturation concentrations are somewhat overestimated at high salt, likely because the model’s implicit, mean-field (Debye-Hückel/Yukawa) treatment of ions omits ion-ion correlations and counterion condensation onto the highly charged chains. These effects strengthen with increasing salt and would otherwise enhance the effective attraction between oppositely charged proteins beyond simple linear screening^65^, thus effectively reducing the saturation concentration to undergo phase-separation. The black curves in Fig. 1b-e are mean-field Flory-Huggins-Voorn-Overbeek fits to the experimental data^66^, shown only as a guide to the eye; all comparisons are made against the measured coexistence concentrations.

For ProT*α*-R50, the *in vitro* data show a re-entrant phase behaviour at high monovalent salt, with a slight decrease in saturation concentration beyond 500 mM. Our model instead predicts condensate formation only up to ~ 500 mM, as the absence of explicit ions precludes the spontaneous recovery of re-entrant phase separation with increasing salt, consistent with previous studies^25^. At low salt concentrations, sampling remains challenging even with coarse-grained simulations, limiting the accuracy of the saturation concentration unless enhanced-sampling or explicit thermodynamic-integration methods are used^67^. For ProT*α*-H1, the model predicts phase separation up to ~170 mM of monovalent salt concentration (e.g., NaCl/KCl), with a dilute-phase saturation concentration of ~5-10 mg/mL between 130 and 170 mM salt concentration (Fig. 1b). This phase diagram provides the thermodynamic reference against which we later interpret the nucleation simulations: at each salt concentration, it sets the protein concentration above which the homogeneous dilute phase becomes supersaturated and a condensate can form.

### Salt-tunable dynamics reproduce the experimental viscoelasticity of complex coacervates

Having established that the model captures the salt-dependent phase behaviour of these complex coacervates, we next test whether it also reproduces the dynamics of the condensed phase, a more stringent benchmark since these emergent properties were not used in its parametrization^48^. To connect molecular-scale dynamics to mesoscale material properties, we quantify three observables, probing increasingly large length scales, for ProT*α* across mixtures and salt concentrations at 295 K (Supplementary Section S4): the conformational reconfiguration time *τ*_*r*_, from the decay of the end-to-end vector autocorrelation function; the translational diffusion time *τ*_*d*_, marking the onset of the Fickian regime in the monomeric mean-squared displacement (MSD); and the condensate viscosity *η*, obtained via the Green-Kubo relation from the time autocorrelation of the pressure tensor^30,68^ and computed with a multiple-*τ* correlator that combines fine resolution at short times with coarser sampling at long times (Fig. 2a-c; see Methods and Supplementary Section S12). Because the reconfiguration time depends on which segment of the sequence defines the correlated vector, we also compute end-to-middle correlations (Fig. 2a): these decay ~25% and ~10% faster than the end-to-end correlations at 50 and 130 mM, respectively, a first signature of the length-scale dependence and also relevant to the interpretation of FRET measurements, which typically probe labelling positions away from the chain termini^15^.

**FIG. 2.**
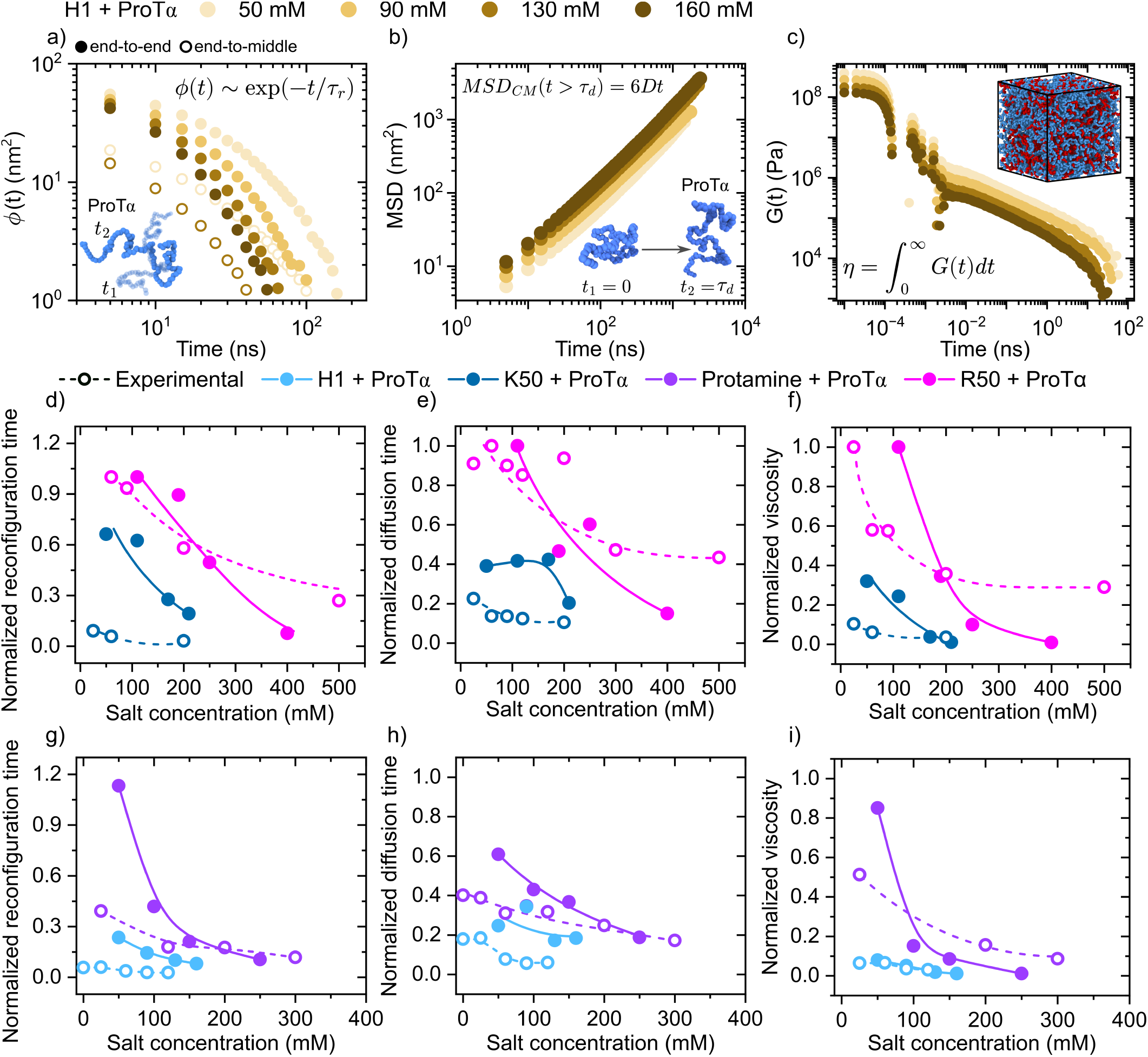
Dynamic transport properties of phase-separated systems from coarse-grained simulations. (a) End-to-end (filled) and end-to-middle (open) vector autocorrelation functions, (b) mean-squared displacement, and (c) stress relaxation moduli of H1+ProT*α* at different salt concentrations. (d-f) Reconfiguration time, diffusion time and viscosity for R50+ProT*α* and K50+ProT*α*; (g-i) the same for protamine+ProT*α* and H1+ProT*α*. Filled symbols/solid lines: simulations; open symbols/dashed lines: experimental data^16^ (both at 295 K). All data normalized by the maximum value for R50+ProT*α*.

When computational predictions and experimental values are normalized by their maximum across the ProT*α*-R50 system (Fig. 2d-i), simulations recapitulate the experimental ordering of *τ*_*r*_, *τ*_*d*_ and *η* across the different mixtures and salt concentrations^16^, despite the implicit-solvent approximation. Deviations are largest at very low salt concentration (i.e., *<* 50 mM), where the terminal relaxation of *G*(*t*) requires extensive simulations beyond ~ 5 *µ*s and the Debye-Hückel approximation begins to break down^42,69^. The achieved near quantitative agreement in normalized viscosity across all four systems indicates that the Mpipi-Recharged captures the coupling between sequence-encoded interactions and emergent material properties^55^, providing a validated basis for the mechanistic analysis of condensate dynamics that follows.

### Length-scale-dependent dynamics decouple fast local intermolecular contacts from slow global relaxation

To identify the physicochemical origin of these dynamics, we next analyse intermolecular contacts and monomeric mobility in ProT*α*-H1 and ProT*α*-protamine condensates at 50 and 160 mM salt (Fig. 3a). Contact maps (Supplementary Sections S8 and S10) show that ProT*α* binding is dominated by acidic hotspots at residues 55-85 and 93-100, formed mostly by glutamic and aspartic acids, with contact number frequencies increasing at low salt due to reduced electrostatic screening^48^, consistent with atomistic simulations^16^. This agreement indicates that the coacervate interaction network is primarily governed by residue-level charge distribution rather than fine atomistic detail. Lysines account for ~60% of H1 contacts and arginines for *>*80% of protamine contacts. Average inter-protein contact lifetimes differ markedly between the two systems: 18.9 ps for ProT*α*-protamine against 8.6 ps for ProT*α*-H1 at 50 mM, and 12.0 against 7.9 ps at 160 mM (Supplementary Fig. S2), an ordering consistent with all-atom simulations of the same coacervates^16^. In both systems the typical contact persists for only a few picoseconds, and the difference resides in the heavy tail of the lifetime probability distribution *P* (*τ*) (Fig. 3b). The two systems also respond differently to screening: protamine contacts shorten by a factor of 1.6 between 50 and 160 mM, whereas H1 contacts change by only 1.1, identifying the arginine-dominated interface as the more strongly electrostatic of the two and tying single-contact stability to the lysine/arginine composition of the contacts themselves (Fig. 3b, right). Representative interaction snapshots illustrate this rapid, local charge-pair switching: in ProT*α*-H1, residues E63-E65 initially pair with K161 of H1 before switching to K158 within ~2 ps timescale, while in ProT*α*-protamine, D95-D96 evolve from an initial D96-R16 pairing towards D95-R16 and D96-R17 on a similar timescale (Fig. 3c; Supplementary Sections S9, S13).

**FIG. 3.**
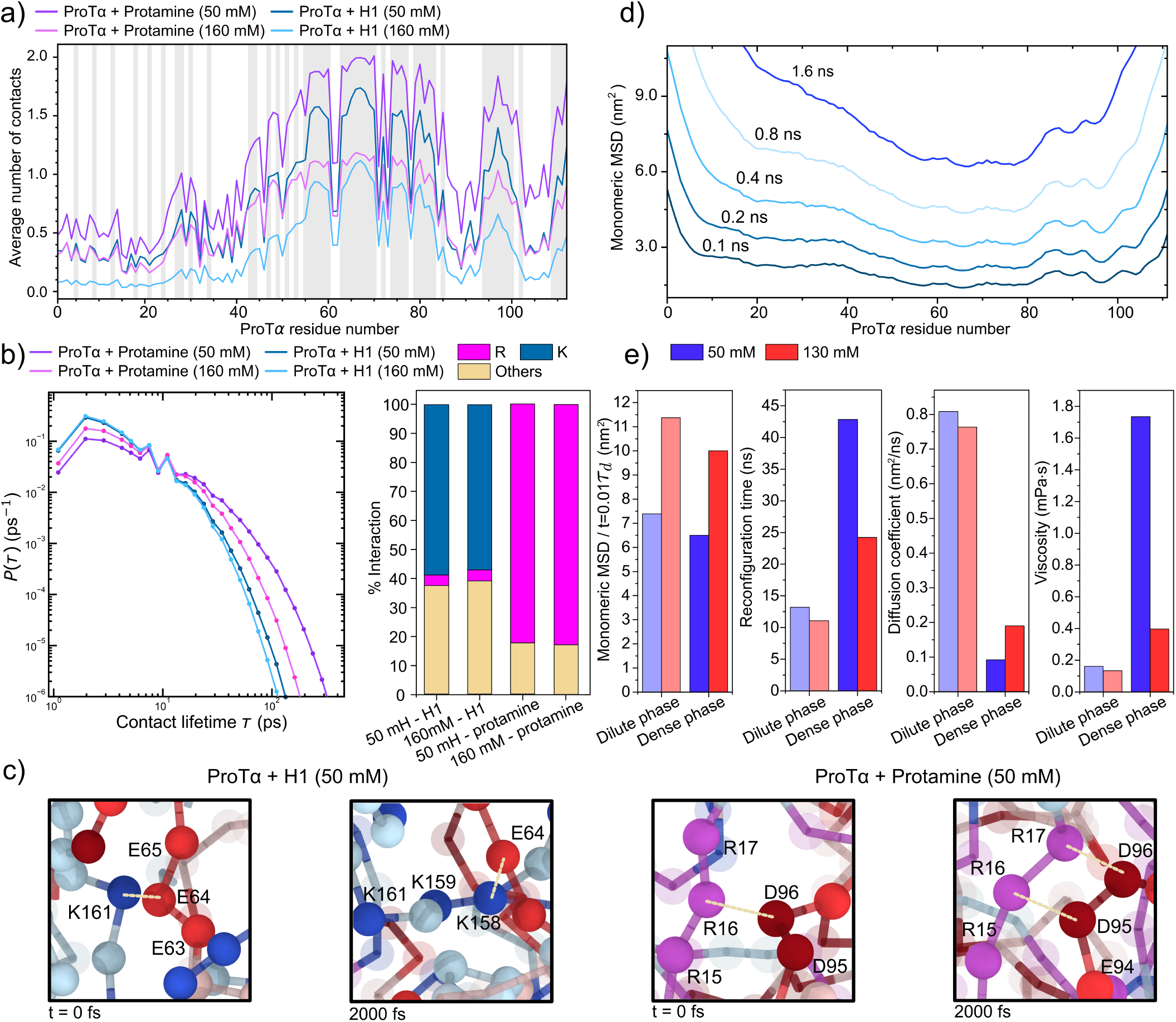
Phase-dependent dynamics and residue-level interactions in ProT*α*-H1 and ProT*α*-protamine systems. (a) Average contacts per ProT*α* residue with H1 (blues) and protamine (purples) at 50 and 160 mM, 295 K; grey shading marks acidic residues. (b) Distribution of inter-protein contact lifetimes *P* (*τ*) for ProT*α*-H1 (dark navy, 50 mM; light blue, 160 mM) and ProT*α*-protamine (purple, 50 mM; magenta, 160 mM) at 295 K, obtained from a transition-based (core-state) contact criterion (Supplementary Section S9); right, breakdown of the fraction of ProT*α* contacts made by lysine (K), arginine (R) and other residues at the same conditions. (c) Snapshots highlighting representative oppositely charged residue pairings at 295 K, 50 mM. (d) Monomeric mean-squared displacement along the ProT*α* sequence at different lag times (H1+ProT*α*, 295 K, 50 mM). (e) Dynamic parameters (monomeric MSD, reconfiguration time, diffusion coefficient, viscosity) compared between the dilute and dense phases of ProT*α*-H1 at 50 and 130 mM, 295 K.

This local reorganization leaves a signature in the monomeric MSD. Unlike a typical homopolymer, whose short-time monomeric MSD is symmetric about the central monomer^70^, the sequence-specific charge patterning of ProT*α* strongly breaks this symmetry (Fig. 3d): residues within persistent-contact regions (60-80, 90-100) show markedly reduced short-time mobility relative to the rest of the chain, reproducing the sequence-dependent profile reported in atomistic simulations^16^ and reinforcing the validity of the coarse-grained description at the residue level (Fig. 3a).

Comparing dynamics in the dilute and dense phases at 50 and 130 mM directly resolves the apparent mismatch between fast local contacts and slow bulk viscosity (Fig. 3e). At short times (*t* = 0.01 *τ*_*d*_), the monomeric MSD of the central ProT*α* residue is comparable in both phases, but *τ*_*r*_, the diffusion coefficient and *η* differ substantially. At 50 mM, chain reconfiguration is ~3-fold faster and diffusion ~ 9-fold faster in the dilute phase than in the condensate (~ 2- and ~ 4-fold, respectively, at 130 mM), and viscosity rises from 0.16 to 1.7 mPa *·*s (50 mM) and from 0.13 to 0.40 mPa*·* s (130 mM) upon condensation. The more dramatic slow-down at 50 mM salt tracks the ~55% increase in condensate density between the two salt conditions (Fig. 1b). These results show that condensation strongly slows large-scale relaxation modes—e.g., viscosity, set by the slowest network components, not by ProT*α* or H1 individually—while leaving sub-Kuhn-scale dynamics essentially unchanged, raising the question of what sets this intermolecular network’s connectivity in the first place.

### Chain entanglement links protein dynamics to condensate viscoelasticity

Because local dynamics are governed by residue-residue interactions with lifetimes orders of magnitude shorter than *τ*_*r*_ or the diffusion time, they can be interpreted within two complementary polymer-physics frameworks for the crossover between unentangled and entangled behaviour^16^: the sticky Rouse model^71,72^, and the Rouse model with entanglements based on Reptation Theory, in which motion becomes confined to a tube imposed by surrounding chains^73–77^. We quantify entanglement directly via primitive-path analysis (PPA;^30,78,79^; Supplementary Section S11) calculations, which reveal a significantly higher density of topological constraints at lower salt concentration (Fig. 4a). Using ⟨ **R**^2^ ⟩ = *a L*_*pp*_, we obtain an average tube diameter *a ≈*7.0 nm and an average entanglement number *Z≈* 1.2-2.5 across conditions (Fig. 4b). As a reference, in the benchmark coarse-grained Kremer-Grest model of polymer melts, chains typically require *N* ≳ 140 beads to reach *Z ~* 1.7 at a monomer density of 0.85^80–83^; the monomer density in our condensates spans 0.27-1.38, placing these systems near the crossover between the Rouse and weakly entangled regimes^82,84^ (Supplementary Section S14).

**FIG. 4.**
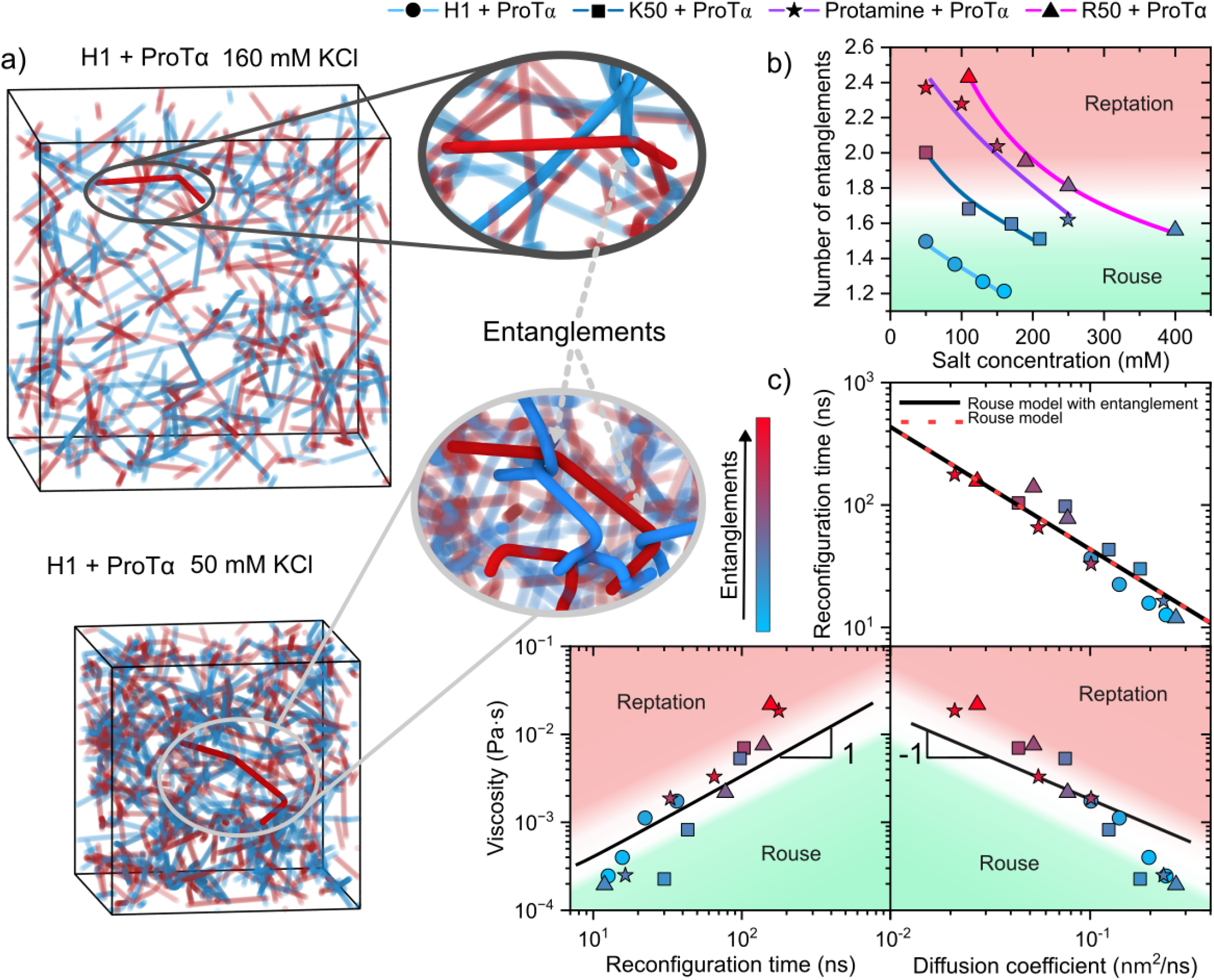
Linking protein dynamics and translational diffusion to condensate viscoelasticity. (a) Primitive-path snapshots of H1 (blue) + ProT*α* (red) at 50 and 160 mM, with entanglements highlighted. (b) Average entanglement number from primitive-path analysis versus salt concentration, with approximate reptation/Rouse regimes shaded. (c) Simulated viscosity, diffusion coefficient and reconfiguration time compared with Rouse and reptation power-law predictions; colour indicates entanglement number (red: more entangled, blue: less entangled).

This crossover has direct material consequences. In the Rouse limit, viscosity scales linearly with chain length and is set by monomeric friction, resulting in relatively low-viscosity condensates. The onset of entanglement, by contrast, introduces topological constraints that dramatically slow protein relaxation, leading to a much steeper scaling (*η ∝ N*^3^ theoretically, *N*^3.4^ experimentally^85,86^)—so that even a modest increase in the number of entanglements per chain can drive a significant change in material response, as seen in the fluid-to-viscous transition of aging protein condensates^18,37,87,88^.

To rationalize this dynamical regime further, we correlate the translational diffusion coefficient *D*, the reconfiguration time *τ*_*r*_, and the condensate viscosity *η*. Since *η* is governed by the terminal relaxation time (*η ~ τ*) while diffusion scales as *D ~* ⟨*R*^2^⟩*/τ*, viscosity and diffusion are expected to be inversely related, *η ~ D*^*−*1^. Across the studied complex coacervates and salt concentrations, the relationship between *τ*_*r*_ and *D* is essentially independent of protein identity, with all systems collapsing onto a single linear trend, while *η* shows two distinct regimes consistent with the entanglement numbers from PPA: systems with *Z* ≳ 2 fall within an entangled regime, whereas less entangled systems follow Rouse-like behaviour (Fig. 4c). Overall, this supports a picture in which condensate dynamics are predominantly Rouse-like, with moderate corrections from transient entanglements depending on the enviromental conditions.

As the system approaches the entangled regime with increasing protein concentrations and interaction strength (*i*.*e*., lower salt), global relaxation slows and viscosity scales far more steeply with chain length^85,86^. Material exchange with the surrounding medium, droplet fusion and shape relaxation become markedly hindered as a result^18,37,89^, showing how small salt-driven changes in network connectivity can trigger a transition from dynamic, liquid-like condensates to kinetically arrested assemblies with impaired material transport^15,16^.

Taken together, Figs. 2-4 establish a length-scale-dependent dynamics of biomolecular condensates. At the residue scale, oppositely charged pairs form and exchange on picosecond timescales, and local mobility is essentially the same in the dilute and dense phases. At the protein scale, reconfiguration, diffusion and condensate viscosity slow down markedly upon self-assembly, by an amount set by the density of the network and by the topological constraints that each biomolecule experiences, both of which grow as salt is reduced in highly charged sequences as those typically present in polyanionic/polycationic complex coacervates. What distinguishes a condensate from the surrounding solution is therefore not the strength or lifetime of individual contacts, but the collective network connectivity that these contacts sustain. This raises the central question of this work: how such a connected network first emerges from a dilute solution of highly charged chains.

### A charge-compensated ProT*α*-H1 dimer is the elementary building block of condensate assembly

Condensate formation from a homogeneous solution is a nucleation process, in which the height of the free energy barrier and the identity of the species that populate the supersaturated metastable solution beforehand set when and where a condensate emerges^39,41,64^. For complex coacervates, the natural candidates for such precritical nuclei formation are the small dynamic complexes that oppositely charged proteins form in the dilute phase. To probe how condensates nucleate from the dilute phase, we first compute potential-of-mean-force (PMF) profiles for ProT*α*-H1 (P, H) complexes at 295 K and 200 mM salt using Adaptive Biasing Force (ABF) simulations^90,91^ (Fig. 5a; Methods; Supplementary Section S5), within a two-state binding model *P* + *H* ⇌ *PH* extended to the trimers PPH and PHH^27^. ABF is well suited to these highly flexible, electrostatically dominated complexes, which lack well-defined free-energy barriers. By adaptively estimating and compensating the mean force along the intermolecular separation, it enhances sampling across bound and unbound states and yields a continuous, well-converged PMF (Fig. 5a). For the PH dimer, the collective variable is the distance between the geometric centres of ProT*α* and H1; for the PPH and PHH trimers, two such distances are used, between one ProT*α*-H1 pair and between the resulting dimer and the third chain.

**FIG. 5.**
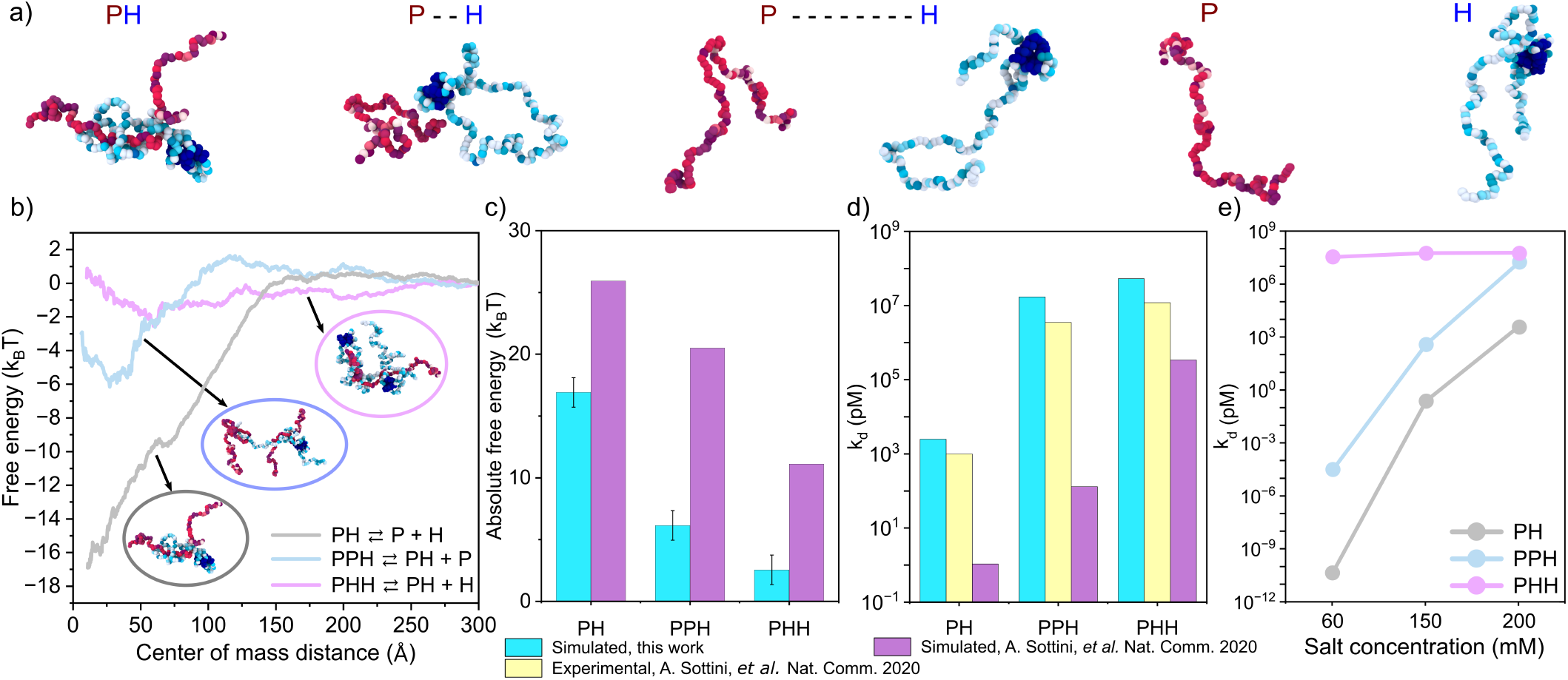
Dilute-phase free-energy landscapes and binding affinities of small ProT*α*-H1 complexes. (a) Representative H1 (H)/ProT*α* (P) configurations along the binding coordinate; the H1 globular domain is highlighted in dark blue. (b) Free-energy profiles for PH (grey), PPH (light blue) and PHH (pink) at 295 K, 200 mM. (c) Comparison with literature PMF values^27^. (d) Dissociation constants: experiment (yellow)^26^, this work (cyan), and literature PMF estimates (purple)^27^. (e) Dissociation constants versus salt (60, 150, 200 mM) for PH, PPH and PHH.

The PH dimer is the most stable species (minimum *≈ −*17*k*_*B*_*T*), consistent with maximal stability near electroneutrality, followed by PPH (*≈ −*6*k*_*B*_*T*) and the markedly weaker PHH (*≈ −*2.5 *k*_*B*_*T*) (Fig. 5b,c), in qualitative agreement with previous CG simulations^26,27^, though Mpipi-Recharged predicts a substantially less stable PPH complex. This hierarchy follows the net charge of each complex: pairing ProT*α* (*Z* = *−*43) with H1 (*Z* = +54) leaves the PH dimer with a net charge of only +11, whereas adding a further ProT*α* or H1 yields complexes with *−*32 (PPH) and +65 (PHH), so that the binding free energy decreases monotonically as uncompensated charge accumulates in the small complex. The resulting dissociation constant for PH, *K*_*d*_ = 2.5 nM, closely matches *in vitro* experimental values (~ 1.0 nM)^26^, and the trimer values (*K*_*d*_ = 17 *µ*M for PPH, 53 *µ*M for PHH) show markedly improved agreement over previous CG models (Fig. 5d)^27^. We attribute this improvement to the pair-specific, asymmetric treatment of attractive and repulsive electrostatics in the Mpipi-Recharged^48^, which departs from the symmetric screened-Coulomb potentials used previously^26^ and is expected to better capture experimental dissociation constants. This effect is amplified in the trimers by their uncompensated charge. Increasing salt weakens PH and PPH binding while leaving PHH largely unaffected, consistent with experiments (Fig. 5e)^26^, an asymmetry attributable to the model’s stronger repulsion between like-charged cationic pairs than between anionic pairs^48^. Together, these results identify the PH dimer as a potential nucleation intermediate from which PPH trimers subsequently assemble, and then grow into mesoscopic post-critical condensates. With a composition close to that of the electroneutral condensate itself, the PH dimer is effectively its minimal charge-compensated unit. Biologically, it corresponds to the ultrahigh-affinity 1:1 complex that ProT*α* forms with the linker histone H1^26^, which we thus identify as the natural precursor of the condensed phase. Directly resolving such rare, transient assembly events experimentally or via atomistic MD remains extremely challenging, whereas CG molecular dynamics gives direct access to the full trajectory of cluster nucleation and growth.

### Brute force simulations reveal a hierarchical, dimer-seeded nucleation pathway

To resolve how these minimal complexes grow into condensate-scale assemblies—the nucleation step linking dilute-phase binding to the phase behaviour of Fig. 1—we perform brute force nucleation simulations of ProT*α*-H1 mixtures starting from homogeneous low-density configurations in the canonical ensemble, so that the stochastic nature of cluster formation is captured directly^92,93^. These simulations span protein densities from 0.0025 g/cm^3^ (comparable to the dilute phase) to 0.04 g/cm^3^ (close to the critical density) and salt concentrations of 60, 150 and 200 mM (see Supplementary Section S15 for further details). Compared against the DC phase diagram (Fig. 1b), these conditions span a full range of supersaturated solutions: at 60 mM, where the experimental saturation concentration lies below 0.1 mg/mL, every density studied is strongly supersaturated; at 150 mM, the lowest densities lie at (or slightly below) the saturation concentration of ~5-10 mg/mL predicted by the model, whereas the highest ones are supersaturated; and at 200 mM, above the highest salt concentration at which the model predicts phase separation (i.e., 170 mM), no condensate is thermodynamically stable at any density or concentration. We first compute the number of protein replicas belonging to clusters of at least 10 proteins (Fig. 6a). At low density, stable clusters form only at 60 mM, where reduced electrostatic screening favours persistent association even at low protein concentration, and clusters grow at a slow but constant pace. In contrast, at 150 and 200 mM no stable clusters are observed, and proteins remain in the dilute phase as free chains or forming transient small complexes. At the highest explored density, all three salt conditions exhibit radically different outcomes: proteins coalesce into a single, near-fully percolated condensate of all 200 chains at 60 mM; fluctuating clusters of 50-100 chains form at 150 mM, remaining in a highly dynamic balance with proteins still free in the dilute phase; and at 200 mM no cluster forms, so spontaneous nucleation is thermodynamically inaccessible (Fig. 6a). Notably, 60 mM is the only salt condition at which nucleation occurs across the two protein densities explored. These outcomes map directly onto the phase diagram: self-assembly is fast and ubiquitous where the solution is strongly supersaturated (60 mM); it is confined to the most concentrated systems close to the phase boundary (150 mM), where small clusters fluctuate strongly and coexist with free proteins in the solution (consistent with the decrease in interfacial tension expected on approaching the critical point^94^ which destabilizes a well-defined droplet); and it is absent in the one-phase region (200 mM).

**FIG. 6.**
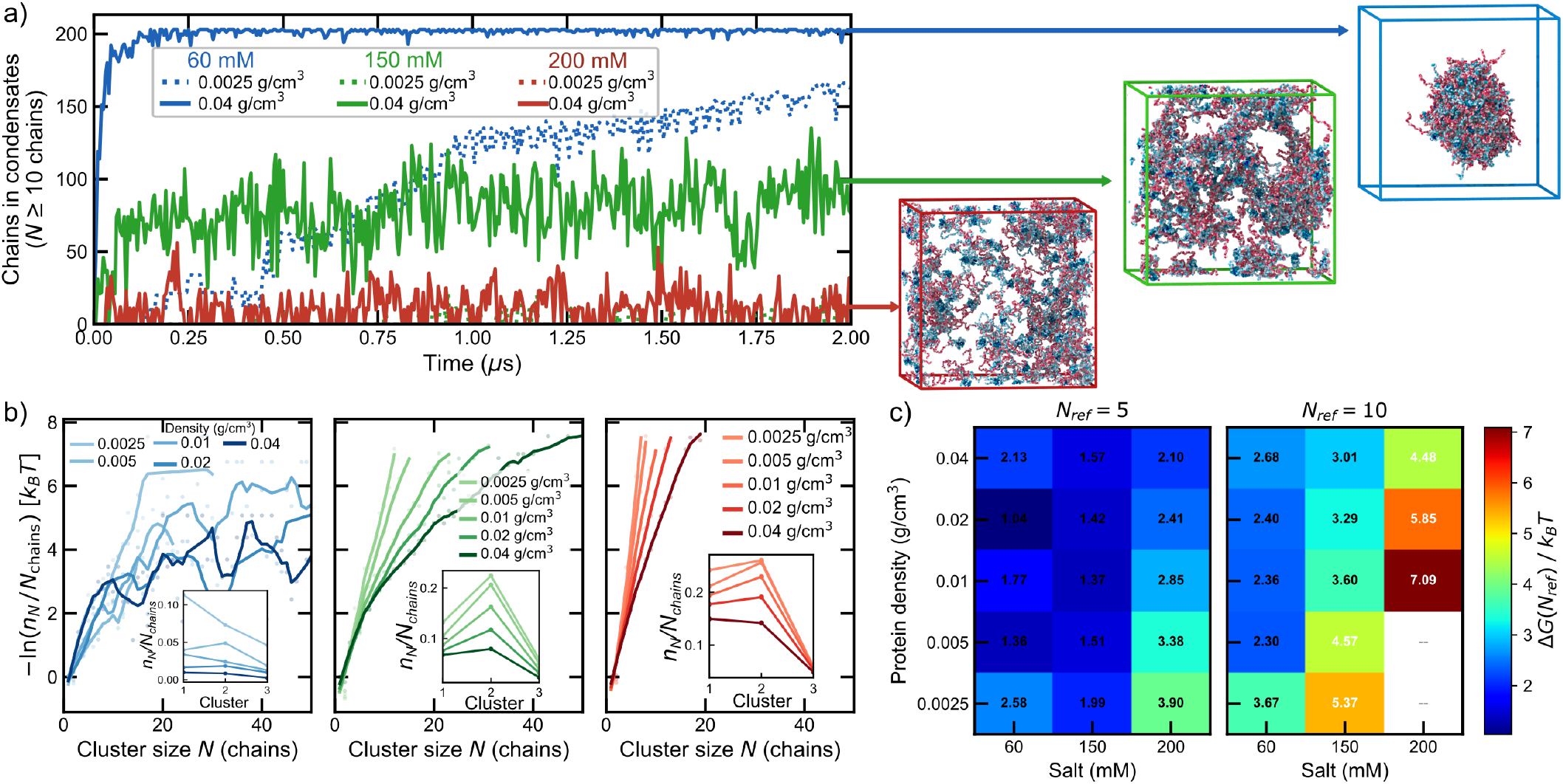
Cluster analysis and nucleation of ProT*α*-H1 condensates. (a) Number of chains condensed into clusters of *N≥* 10 chains versus simulation time (2 *µ*s) at 60 (blue), 150 (green) and 200 mM (red) salt, for the most dilute (0.0025 g/cm^3^, dotted) and most concentrated (0.04 g/cm^3^, solid) densities simulated, with representative condensate snapshots at 2 *µ*s (ProT*α* pink, H1 blue), boxed in the corresponding salt colour. (b) Cluster-size distribution, *−* ln(*n*_*N*_ */N*_chains_), normalized by total chain number for systems at 60, 150 and 200 mM salt concentration (light-to-dark shading: protein density from 0.0025 to 0.04 g/cm^3^). Insets show *n*_*N*_ */N*_chains_ versus cluster size, highlighting the dimer-peaked pre-nucleation population of the metastable phase. (c) Nucleation free energy barrier Δ*G*(*N*_ref_) at a fixed, well-sampled reference cluster size (*N*_ref_ = 5, left; *N*_ref_ = 10, right) versus salt concentration and protein density; blank cells mark conditions where *N*_ref_ is too rarely sampled for a reliable estimate.

We next compute the nucleation free energy barrier as a function of cluster size, Δ*G*(*N*) = *−*ln(*n*_*N*_ */N*_chains_), which can be obtained from the population *n*_*N*_ of clusters of size *N* sampled across the metastable pre-nucleation time of each trajectory^39^ (Supplementary Section S15). This barrier is visibly noisy at 60 mM (Fig. 6b), where the system nucleates into a single, near-fully percolated cluster so rapidly that few genuinely pre-nucleation configurations remain to populate the intermediate cluster sizes. Despite this scatter, the barrier increases consistently with higher salt concentration (green and red panels) across the cluster sizes that are well sampled at all three conditions, an ordering that sharpens markedly for larger clusters. Within classical nucleation theory (CNT) framework, Δ*G*(*N*) results from the competition between the free energy gain of transferring *N* chains into the dense phase, *−N* Δ*µ*, and the interfacial penalty, *∝ γN*^2/341^. At fixed density, increasing salt raises the saturation concentration (Fig. 1b) and therefore lowers the effective supersaturation and Δ*µ*. That Δ*G*(*N*) nonetheless rises with salt, despite the decrease in interfacial tension expected on approaching the critical salt concentration, indicates that the loss of supersaturation dominates; this also explains why the salt ordering becomes more pronounced for larger clusters, for which the bulk term carries more weight on the nucleation barrier. The pre-critical cluster population (inset plots in Fig. 6b) peaks at the dimer at 150 and 200 mM, and under all conditions the dimer is more populated than the trimer or higher-order complexes (at 60 mM the peak is not resolved because nucleation is too fast to sample the metastable state; Supplementary Section S15). This identifies the PH dimer as the effective seed for nucleation, in direct correspondence with its markedly higher stability relative to PPH and PHH in the dissociation-constant hierarchy established above (Fig. 5b). Nucleation of ProT*α*-H1 condensates is therefore hierarchical: chains first pair into charge-compensated PH dimers, which dominate the metastable supersaturated solution before condensates emerge, and it is the subsequent association of these dimers—and of the much weaker trimers—into larger clusters that carries the nucleation barrier. This two-level pathway parallels the role of ion pairs and small ionic clusters in the nucleation of crystals from supersaturated electrolyte solutions^39,95^, with the difference that here the elementary “ion pair” is itself a highly disordered, dynamic complex of two polyelectrolytes.

We now evaluate at two fixed, well-sampled reference sizes (*N*_ref_ = 5 and 10; Fig. 6c) the barrier’s sensitivity to salt and density changes with cluster size. For *N*_ref_ = 5 it is comparable across densities at 60 and 150 mM (*≈*1-2.6 *k*_*B*_*T*) and rises above this range specifically at 200 mM and low protein concentrations, where it is highest (*≈*4 *k*_*B*_*T*). For *N*_ref_ = 10, density becomes far more important, with the barrier staying low across the full density range at 60 mM, increasing substantially at 150 mM, and becoming largest at 200 mM, where forming a cluster of this size is so rare at the lowest densities that a reliable initial part of the free energy barrier cannot be estimated (blank cells, Fig. 6c). At 150 mM, the barrier for *N*_ref_ = 10 drops from 5.4 to 3.6 *k*_*B*_*T* as the density increases from 0.0025 to 0.01 g/cm^3^ and then levels off (3.3 and 3.0 *k*_*B*_*T* at 0.02 and 0.04 g/cm^3^ respectively). Its steepest variation occurs as the simulation protein density crosses the saturation concentration predicted by DC at this salt (~0.005-0.01 g/cm^3^; Fig. 1b), an independent consistency check between the unbiased brute force simulations and the Direct Coexistence simulations. This size-dependent cluster crossover, from a barrier that is comparatively insensitive to density at small *N* to one that is strongly density-dependent at larger *N*, marks the transition from initial dimer/trimer formation to larger pre-critical condensate-scale growth. The smallest clusters are essentially the complexes characterized in Fig. 5, whose stability is set by pairwise electrostatic binding and hence by screening, whereas larger clusters require many protein to meet, and therefore probe the supersaturation of the solution. Together with the PMF results, this indicates that ProT*α*-H1 nucleation is governed by a salt-dependent balance between the intrinsic stability of the PH dimer and the collective energetics of larger-scale assembly, rather than by a single critical cluster size, and establishes the hierarchical PH-dimer-seeded pathway as the route by which the condensates characterized in the preceding sections actually form.

Salt therefore acts at every level of the hierarchy resolved in this work. It sets the stability of the elementary PH complex (Fig. 5), the supersaturation that drives the collective assembly of these complexes into condensate-scale clusters (Figs. 1 and 6), and, once the condensate has formed, the density, connectivity and entanglement that control its material properties (Figs. 2-4). The same reduction in electrostatic screening that lowers the nucleation barrier also yields denser, more entangled and more viscous condensates, so that the conditions that favour condensate formation also favour a less dynamic mature state. Biologically, ProT*α*-H1 at physiological ionic strength (~ 150 mM) lies close to the boundary between a solution dominated by soluble, high-affinity PH complexes and one that nucleates condensates only at moderately high protein concentrations; such proximity to the nucleation threshold would allow condensate formation to be switched on or off by modest changes in local concentration or ionic environment, rather than by changes in the underlying intermolecular interactions.

## DISCUSSION

In this work, we show that ProT*α*-H1 complex coacervates nucleate via a two-step, dimer-seeded pathway: oppositely charged proteins first pair into a charge-compensated dimer in the dilute phase, and it is the subsequent, concentration- and salt-dependent assembly of these dimers that carries the nucleation barrier toward mesoscopic stable condensates. We resolve this pathway using a residue-level, coarse-grained description of these complex coacervates that we validate across their phase diagram, protein dynamics and material properties, recapitulating the salt- and sequence-dependent stability, binding affinities and viscosities measured *in vitro* for ProT*α*-based condensates^15,16^. This agreement indicates that our model captures the key electrostatic interactions driving phase separation and provides a ro-bust thermodynamic framework for understanding condensate formation at submolecular level, and how these properties are primarily governed by the sequence distribution of charged residues rather than by fine atomistic detail. Microscopic heterogeneity along the protein sequences gives rise to a network of transient interactions that couples rapid local amino acid rearrangements to slower internal protein reconfiguration and translational diffusion at condensate level.

Within the condensate, our results reveal an intrinsically length-scale-dependent picture of protein dynamics: short-lived, labile residue-residue contacts rearrange rapidly, and equally fast, in the dilute and dense phases, yet coexist with global relaxation modes—chain reconfiguration and translational diffusion—that slow down markedly upon condensation, producing the highly viscous character of biomolecular condensates. Resolving this apparent contradiction requires probing the relevant relaxation timescales at different length scales, which are difficult to access without coarse-grained simulations^30,53^. Condensate viscosity reflects the collective relaxation of the inter-protein network rather than the local mobility of individual residues, so that condensate formation strongly alters protein diffusion and reconfiguration while sub-Kuhn-scale dynamics at residue-level remain largely unaffected between phases. Primitive-path analysis places these systems near the Rouse-to-entangled crossover, with reptation effects becoming significant only at the highest densities (i.e., lowest salt concentration), directly linking the emergence of topological constraints to the transition from liquid-like to kinetically arrested condensates—a transition with direct functional consequences, since it hinders material exchange with the surrounding medium and processes such as droplet fusion and shape relaxation^21,37,87,88^. What sets a condensate apart from the surrounding solution is therefore not the lifetime of individual residue-residue contacts, but the connectivity of the network they sustain.

We elucidate how that network first assembles across the condensate nucleation process. Dilute-phase binding calculations identify the ProT*α*-H1 dimer as the minimal, most stable building block, driven by strong electrostatic complementarity and consistent with the experimental hierarchy of binding affinities, while higher-order trimers become progressively less stable as they accumulate uncompensated charge (Fig. 5). Crucially, the dimer dominates the metastable pre-critical cluster population in unbiased simulations, confirming a hierarchy first established thermodynamically from the dissociation constants (Fig. 6). Brute-force nucleation simulations confirm that these dimers act as the minimal nucleation seeds from which larger, condensate-scale clusters subsequently grow, and show that both the onset of clustering and the associated nucleation barrier are jointly set by protein and salt concentration, rather than by a single-parameter critical nucleus size. The barrier consistently increases at higher salt concentrations due to dominant electrostatic screening which disables collective growth at the salt concentration of *≈*200 mM. This outcome is fully consistent with the independently computed phase diagram: nucleation is spontaneous where the solution is strongly supersaturated, confined to the most concentrated systems close to the phase boundary, and absent in the one-phase region. Since the trajectories begin from homogeneous, unseeded configurations, they capture this stochastic assembly process directly rather than presupposing a nucleus shape or critical size. To our knowledge, this is the first time spontaneous, unseeded nucleation has been resolved for highly charged protein complex coacervates. From a physical standpoint, the condensation of these proteins therefore follows a dimer-driven nucleation pathway, in which a charge-compensated elementary unit—a highly disordered, dynamic protein dimer—forms first, and the nucleation barrier is carried by the collective self-assembly of these units. The hierarchical, dimer-seeded pathway thus provides the missing link between dilute-phase binding, the thermodynamic stability established by the phase diagrams and the mesoscale material properties characterized once the condensate has formed, and situates nucleation, rather than the mature condensate alone, as the defining stage of the interaction dynamic network underlying complex coacervation.

Together, this work establishes a predictive framework which connects sequence-encoded intermolecular forces to condensate dynamics, material properties and nucleation, spanning from the earliest stages of dimer-mediated nucleation to the emergent viscoelastic properties of the mature condensates. It further demonstrates that residue-level coarse-grained models can reach spatio-temporal regimes often inaccessible to atomistic or slab-based simulations with a certain degree of accuracy^45^. Notably, electrostatic screening governs both ends of this pathway—the barrier a condensate must cross to be formed, and the entanglement and viscosity it acquires once formed—a coupling with direct implications for how condensate function is tuned *in vivo* and engineered *in vitro*^18,21,96^. Since the stability of the elementary building block follows the charge balance between partners, subtle changes in local ionic strength or protein concentration, as well as modifications that alter the charge of either partner, such as phosphorylation or lysine acetylation, would be expected to shift where and when these condensates nucleate, offering physically-grounded routes by which cells could regulate condensate formation and their viscoelastic behaviour.

## METHODS

### Model and simulations

All simulations use the residue-level coarse-grained Mpipi-Recharged force field^48^, an implicit-solvent, implicit-ion model with pair-specific Wang-Frenkel short-range and Yukawa electrostatic interactions (Supplementary Section S1). Direct Coexistence phase diagrams, system compositions and general NVT simulation protocols (LAMMPS^97^, Nosé-Hoover thermostat for rigid globular domains, Langevin thermostat for flexible regions, 10 fs timestep) are detailed in Supplementary Sections S2-S4.

### Dynamic observables

The reconfiguration time *τ_r_* is obtained from a double-exponential fit to the end-to-end vector autocorrelation function; the diffusion time *τ*_*d*_ is defined as the time at which ≥90% of monomeric MSDs reach an apparent exponent *α ≥* 0.9; viscosity is obtained from the Green-Kubo integral of the pressure-tensor autocorrelation function, fitted to a sum of Maxwell modes using RepTate^98^. Time correlation functions are computed with a multiple-*τ* correlator^99,100^. Full definitions and dilute-phase viscosity estimation via the Stokes-Einstein relation are given in Supplementary Sections S6-S7.

### Contact analysis

Static intermolecular contact maps use a sequence-dependent cutoff of 1.2*σ*_*ij*_. Contact lifetimes instead use a transition-based (core-state) criterion, tracked frame-by-frame between residue pairs: a contact forms once the separation first drops below 1.2*σ*_*ij*_ and is taken to persist, irrespective of brief excursions, until it exceeds an outer threshold of 1.8*σ*_*ij*_, so that a recorded rupture reflects genuine unbinding rather than thermal recrossing of a single cutoff (Supplementary Sections S8-S9).

### Primitive-path analysis

Chain entanglement is quantified following the algorithm of Sukumaran *et al*.^78^, minimizing chain contour length with fixed chain ends (Supplementary Section S11).

### Free-energy and nucleation simulations

Binding PMFs use the Adaptive Biasing Force method^90,91^ in Colvars^101^; dissociation constants follow 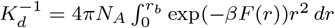 (Supplementary Section S5). Brute-force nucleation simulations of 200 chains were run for 2 *µ*s in the NVT ensemble at protein densities of 0.0025-0.04 g/cm^3^ and 60/150/200 mM salt, with chains assigned to the same cluster below a 50 Åcentre-of-mass separation.

## ACKNOWLEDGEMENTS

The authors gratefully acknowledge conversations with Jeetain Mittal and Ben Schuler which inspired this research. We also thank Nicola Galvanetto for providing the experimental data and for insightful discussions regarding their work. E.P. acknowledges funding from the European Social Fund Plus and project PID2022-136919NA-C33 from the Spanish MICIU. A.R.T. acknowledges funding from MICIU under the Juan de la Cierva fellowship (JDC2024-053759-I). D.d.S. acknowledges funding from MICIU/AEI/10.13039/501100011033 and “ERDF A way of making Europe” through grant PID2024-158678NB-I00. R.C.-G. acknowledges funding from UKRI EPSRC [EP/Z002028/1], following funding from the ERC Consolidator Grant “ChromatinDroplets” under the European Union’s Horizon Europe programme. P.L. acknowledges funding from the European Union’s Horizon 2020 programme (grant agreement 101160499 to J.R.E). J.R.E. acknowledges funding from Emmanuel College, University of Cambridge, the Ramon y Cajal fellowship (RYC2021-030937-I), Spanish grant PID2022-136919NA-C33, and the ERC under Horizon Europe (grant agreement no. 101160499). The authors thank RES for computational resources provided by Mare Nostrum 5 (activity FI-2025-3-0065).

## AUTHOR CONTRIBUTIONS

