## Supplementary Material for "A Dimer-Seeded Pathway Governs the Nucleation of ProT*α*-H1 Biomolecular Condensates"

### Supplementary Information for: A Dimer-Seeded Pathway Governs the Nucleation of ProT$\alpha$-H1 Biomolecular Condensates

Eduardo Pedraza[†,1,2] Andrés R. Tejedor[†,1,2] Álvaro S. Zorita,[1] Rosana Collepardo-Guevara,[3,4,5] David de Sancho[c,6,7] Pablo Llombart [b,1] and Jorge R. Espinosa[a1,2,4,5]

[1]*Department of Physical Chemistry,*
*Universidad Complutense de Madrid,*
*Av. Complutense s/n, Madrid 28040, Spain*

[2]*Instituto Pluridisciplinar, Universidad Complutense de Madrid,*
*P.º de Juan XXIII, 1, Moncloa - Aravaca, 28040 Madrid, Spain*

[3]*Department of Genetics, University of Cambridge,*
*Downing Street, Cambridge CB2 3EH, UK*

[4]*Yusuf Hamied Department of Chemistry, University of Cambridge,*
*Lensfield Road, Cambridge CB2 1EW, UK*

[5]*PhAsIca Biosciences S.L, Calle Velázquez, 27, 28001 Madrid, Spain*

[6]*Polimero eta Material Aurreratuak: Fisika,*
*Kimika eta Teknologia, P Manuel Lardizabal 3,*
*Donostia/San Sebastian 20018, Spain*

[7]*Donostia International Physics Center (DIPC),*
*P Manuel Lardizabal 3, Donostia/San Sebastian 20018, Spain*

(Dated: October 2, 2026)

[a]

[b]

[c]

[†]These authors contributed equally

#### S1. THE MPIPI-RECHARGED MODEL

The Mpipi-Recharged model is a residue-level coarse-grained force field for both protein and protein/RNA condensates [1]. Each amino acid or nucleotide is represented by a single bead connected to adjacent residues by harmonic bonds. Globular domains of the proteins are treated as rigid bodies whose beads are fixed at the $C_\alpha$ positions of the corresponding experimental (Protein Data Bank) or AlphaFold-predicted structure. The potential energy is computed as the sum of pairwise bonded ($E_{\text{bonded}}$) and non-bonded ($E_{\text{non-bonded}}$) interactions as:

$$E = E_{\text{bonded}} + E_{\text{non-bonded}}. \tag{S1}$$

The intrinsically disordered regions (IDRs) aremodelled as fully flexible polymers and these are connected to the globular domains. The bonded potential is written as:

$$E_{\text{bonded}}(r_{ij}) = \sum_{ij} k(r_{ij} - r_0)^2, \tag{S2}$$

where $r_{ij}$ is the distance between the connected beads, $r_0 = 3.81\,\text{Å}$ and $r_0 = 5.00\,\text{Å}$ are the equilibrium bond lengths for protein and RNA, respectively. The spring constant $k = 9.6\,\text{kcal}\cdot\text{mol}^{-1}\cdot\text{Å}^{-2}$. The sum runs over all bonded residues.

Non-bonded interactions consist of the sum of the hydrophobic interaction and the electrostatic interaction. The hydrophobic interaction is given by the Wang–Frenkel (WF) potential [2] that accounts for short-ranged excluded-volume repulsion and long-ranged attraction. This potential is defined as:

$$E_{\text{WF}}(r_{i,j}) = \sum_{ij} \epsilon_{ij}\alpha_{ij}\left[\left(\frac{\sigma_{ij}}{r_{ij}}\right)^{2\mu_{ij}} - 1\right]\left[\left(\frac{R_{ij}}{r_{ij}}\right)^{2\mu_{ij}} - 1\right]^{2\nu_{ij}}, \tag{S3}$$

where

$$\alpha_{ij} = 2\nu_{ij}\left(\frac{R_{ij}}{\sigma_{ij}}\right)^{2\mu_{ij}}\left\{\frac{2\nu_{ij}+1}{2\nu_{ij}\left[\left(\frac{R_{ij}}{\sigma_{ij}}\right)^{2\mu_{ij}} - 1\right]}\right\}^{2\nu_{ij}+1}. \tag{S4}$$

Here $\sigma_{ij}$ is the pair-of-beads diameter, defined from the individual diameters ($\sigma_i$ and $\sigma_j$) assuming the Lorentz-Berthelot mixing rules (*i.e.*, $\sigma_{ij} = (\sigma_i + \sigma_j)/2$). $R_{ij} = 3\sigma_{ij}$ is the cut-off distance for the $ij$-th interaction. The interaction parameter $\epsilon_{ij}$ is defined for each specific amino acid pair based on our atomistic Potential of Mean Force calculations and bioinformatics data [1]. The exponent $\nu_{ij}$ is set to 1 for all pairs and $\mu_{ij}$ depends on the specific pair, ranging from 2 to 12 (see Ref. [1]). Notice that higher values of $\mu_{ij}$ lead to a steeper increase in the repulsive part of the potential. The interaction involving globular domains are reduced to account for the 'buried' interactions. In particular, the interaction between flexible regions and globular domains are screened by a factor of $\sqrt{0.7}$ and the WF interaction between residues in globular regions is scaled down a factor of 0.7.

Electrostatic interactions are described by the Yukawa potential [3] instead of the Debye–Hückel potential [4] originally employed in the Mpipi model. Avoiding the use of explicit charge values, this potential allows to modulate independently the strength of electrostatic interactions in a pair-specific basis. This potential is defined as:

$$E_{\text{electrostatic}} = \sum_{ij} \frac{A_{ij}}{r_{ij}} \exp(-\kappa r_{ij}), \qquad r_{ij} < r_c, \tag{S5}$$

where $A_{ij}$ is the interaction parameter, $\kappa$ is the screening due to ions, and $r_{ij}$ is the distance between interacting residues. The cut-off for the electrostatic interaction, $r_c$, is set to 3.5 nm. The screening parameter $\kappa$ is expressed in a explicit way as $\kappa = \sqrt{8\pi B c_s}$, where $c_s$ is the salt concentration—normally set to 150 mM monovalent electrolyte—and $B = e_0^2/4\pi k_B T \varepsilon_0 \varepsilon_r$ is the Bjerrum length. The relative dielectric constant $\varepsilon_r$ varies with temperature according to the empiric formula [5]:

$$\varepsilon_r(T) = \frac{5321}{T} + 233.760 - 0.9297T + 1.417 \cdot 10^{-3} T^2 - 8.292 \cdot 10^{-7} T^3, \tag{S6}$$

for $T$ in Kelvin. The pair-specific optimized values of $A_{ij}$ of the Yukawa potential can be found in Ref. [1]. In the Mpipi-Recharged model, such parameters indicate that the interaction between oppositely charged pairs is significantly stronger than those between identically charged pairs.

#### S2. SEQUENCES AND PROTEIN COMPOSITIONS OF THE SIMULATED SYSTEMS

##### ProT$\alpha$

GPMSDAAVDTSSEITTKDLKEKKEVVEEAENGRDAPANGNANEENGEQEADNEVDEEEEEGGEEE
EEEEEGDGEEEDGDEDEEAESATGKRAAEDDEDDDVDTKKQKTDEDD

**H1**

MTENSTSAPAAKPKRAKASKKSTDHPKYSDMIVAAIQAEKNRAGSSRQSIQKYIKSHYKVGENADS
QIKLSIKRLVTTGVLKQTKGVGASGSFRLAKSDEPKKSVAFKKTKKEIKKVATPKKASKPKKAASK
APTKKPKATPVKKAKKKLAATPKKAKKPKTVKAKPVKASKPKKAKPVKPKAKSSAKRAGKKK

The globular region of H1 (residues 28-38, 47-57, 64-78, 81-85 and 92-95) is taken from the AlphaFold-predicted structure [6, 7] of UniProt entry P07305 and simulated as a rigid body.

**K50**

KKKKKKKKKKKKKKKKKKKKKKKKKKKKKKKKKKKKKKKKKKKKKKKKKK

**R50**

RRRRRRRRRRRRRRRRRRRRRRRRRRRRRRRRRRRRRRRRRRRRRRRRRR

**Protamine**

MPRRRRSSSRPVRRRRRPRVSRRRRRRGGRRRR

**System composition**

The simulated systems consisted of mixtures of ProT$\alpha$ and a second component. The number of molecules of each species in the simulation box depended on the mixture considered. The compositions are summarized in the following table:

| System | ProT$\alpha$ | Second component |
|---|---|---|
| ProT$\alpha$ + H1 | 113 | 90 |
| ProT$\alpha$ + R50 | 116 | 100 |
| ProT$\alpha$ + K50 | 116 | 100 |
| ProT$\alpha$ + Protamine | 88 | 180 |

**TABLE S1:** Number of molecules of each component in the simulated systems.

#### S3. CALCULATION OF THE PHASE DIAGRAMS VIA DIRECT COEXISTENCE SIMULATIONS

The determination of the phase diagrams in the salt-density plane at 295 K was performed using the Direct Coexistence method [8, 9]. The proteins are placed in a prismatic elongated box to simulate both the high-density and low-density phases separated by an interface. The long side of the box is perpendicular to the interfaces. Since determination of the optimal box dimensions is key to minimizing finite size effects [10] while ensuring computer efficiency, we provide the following guidelines

1. We choose the number of molecules of each component depending on the specific protein mixture to ensure stable phase coexistence. The compositions used in the direct coexistence simulations are reported in Table S1.

2. We enforce the short sides of the slab box to be larger than at least twice the radius of gyration of the protein in the system to avoid self-interactions through the periodic boundary conditions.

3. The long side of the slab box should keep the total density of the system at approximately $\sim$0.1 g·cm$^{-3}$.

Simulations are carried out in the canonical (NVT) ensemble using a Nosé-Hoover thermostat [11] for the rigid bodies (representing the globular domains) included in the RIGID package, and a Langevin thermostat [12] for the particles in flexible regions, both with a relaxation time of 5 ps. The time step for the Verlet integration of the equations of motion is 10 fs. After an equilibration period ($\sim$ 10ns), we run the simulation for 1-3 $\mu$s depending on the specific system. If two different phases are detectable—*i.e.*, a high density phase and a low density phase—the densities are calculated.

#### S4. SIMULATION DETAILS

All quantities reported in this work were computed from NVT simulations carried out at the coexistence densities extracted from the salt-density phase diagrams previously determined by direct coexistence (see Section S3). The system sizes were chosen accordingly to represent the corresponding coexisting phases. All NVT simulations were carried out using the molecular dynamics LAMMPS software [13], version 2nd of August of 2023. We use a Nosé-Hoover thermostat [11] for the rigid bodies (representing the globular domains) included in the RIGID package, and a Langevin thermostat [12] for the particles in flexible regions, both with a relaxation time of 5 ps. The time step for the Verlet integration of the equations of motion is 10 fs. After equilibration has been reached, we run 3-5 $\mu$s production simulations. The cut-off values used for the interactions are those described in Section S1.

#### S5. CALCULATION OF PMFS AND $K_d$ EVALUATION

Enhanced sampling molecular dynamics simulations were performed using the Adaptive Biasing Force [14, 15] (ABF) method as implemented in the Colvars module [16]. The collective variable (CV) was defined as the distance between the geometric centres of selected atomic groups corresponding to different oligomeric subunits. In all cases, a bin width of 0.2 Åwas employed, with lower and upper boundaries set to 0 and 300 Å, respectively. A harmonic upper wall restraint with a force constant of 5.0 kcal $\cdot$ mol$^{-1}$ $\cdot$ Å$^{-2}$ was applied to limit the sampling range where indicated. The minimum number of full samples required before applying the ABF was set to 500 per bin, and the Jacobian term was explicitly included. To quantify the affinity between ProT$\alpha$ and H1 chains at equilibrium, we calculate the dissociation constants $K_d$ from the free energy profiles as $K_d^{-1} = 4\pi N_A \int_0^{r_b} \exp(-\beta F(r)) r^2 \, dr$, where $N_A$ is Avogadro's number, $\beta = 1/(k_B T)$, and $r_b$ delimits the bound-state domain, taken as the separation at which $F(r)$ reaches the plateau of the unbound state or, when present, the maximum of the free-energy barrier separating bound and unbound configurations.

#### S6. EVALUATION OF THE VISCOSITY

In an isotropic system, the Green-Kubo relation allow to obtain the viscosity, $\eta$, by integrating the shear relaxation modulus $G(t)$, *i.e.*, $\eta = \int_0^\infty G(t)dt$. $G(t)$ can be calculated after evaluating the components of the pressure tensor ($\sigma_{\alpha\beta}$) as arising from the virial theorem, as follows [17]:

$$G(t) = \frac{V}{5k_BT}\left[\langle\sigma_{xy}(0)\sigma_{xy}(t)\rangle + \langle\sigma_{xz}(0)\sigma_{xz}(t)\rangle + \langle\sigma_{yz}(0)\sigma_{yz}(t)\rangle\right] \\ + \frac{V}{30k_BT}\left[\langle N_{xy}(0)N_{xy}(t)\rangle + \langle N_{xz}(0)N_{xz}(t)\rangle + \langle N_{yz}(0)N_{yz}(t)\rangle\right], \tag{S7}$$

where $N_{\alpha\beta} = \sigma_{\alpha\alpha} - \sigma_{\beta\beta}$ is the normal stress difference. Finally, $\eta$ is obtained following a two-step procedure. First, we numerically integrate the first part of the decay of $G(t)$ up to $t_0$, thus obtaining the intramolecular contribution to the viscosity ($\eta_0$). Second, in the long time limit, the slower relaxation is integrated by fitting $G(t)$ to the sum of $m$=4–6 Maxwell modes, depending on the span and noise of $G(t)$. The discrete Maxwell modes have the exponential form $G_i \exp(-t/\tau_i)$ and are equidistant in logarithmic time [18]. The fit is carried out with the help of the open-source RepTate software [19]. Finally, viscosity is obtained by adding the two terms:

$$\eta = \eta_0 + \sum_{i=0}^{m} \tau_i G_i \exp\Big(-t/\tau_i\Big), \tag{S8}$$

where $\tau_i$ and $G_i$ are the fitting parameters of $G(t)$ written as the sum of Maxwell modes. Typically, simulations of 3-5 $\mu$s are needed to observe a clear terminal decay in $G(t)$, characteristic of liquid phases. Storage ($G'$) and loss ($G''$) moduli can be directly obtained as the Fourier transform of $G(t)$ also using RepTate [19].

The components of the pressure tensor were evaluated in bulk conditions at the dense-phase coexistence densities (Section S4) in the NVT ensemble employing a Nosé-Hoover thermostat [11] for the rigid bodies (representing the globular domains) integrated in the RIGID package, and a Langevin thermostat [12] for the particles in flexible regions, both with a relaxation time of 5 ps and a time step for the Verlet algorithm of 10 fs. The configurations were generated by placing the protein replicas indicated in Section S2 for each component in a cubic box.

**Dilute-phase viscosity estimation**

The viscosity of the dilute phase was estimated from NVT simulations of a ProT$\alpha$–H1 dimer using the Stokes–Einstein relation [20]:

$$D = \frac{k_BT}{6\pi\eta R_h}, \tag{S9}$$

where $D$ is the diffusion coefficient, $k_B$ is the Boltzmann constant, $T$ the temperature, and $R_h$ the hydrodynamic radius of the protein complex. The hydrodynamic radius, $R_h$, was computed from NVT trajectories using the Kirkwood relation:

$$\frac{1}{R_h} \approx \frac{1}{N^2} \sum_{i=1}^{N} \sum_{\substack{j=1 \\ j \neq i}}^{N} \left\langle \frac{1}{r_{ij}} \right\rangle , \tag{S10}$$

where $N$ is the total number of residues in the dimer and $r_{ij}$ is the distance between residues $i$ and $j$.

#### S7. CALCULATION OF DIFFUSION COEFFICIENTS DIFFUSION TIMES AND RECONFIGURATION TIME

The diffusion coefficient of the proteins inside the condensates is obtained from the mean squared displacement (MSD) of the centre of mass of each set of proteins. After a subdiffusive regime (*i.e.*, ∼1 molecular diameter), proteins exhibit a diffusive behaviour and, therefore, the MSD of the centre of mass can be measured as:

$$\left\langle (\mathbf{R}_{\mathrm{CM}}(t) - \mathbf{R}_{\mathrm{CM}}(0))^2 \right\rangle = 6Dt^{\alpha} \tag{S11}$$

where $\mathbf{R}_{\mathrm{CM}}$ indicates the centre of mass of a given protein at different times, $D$ corresponds to the diffusion coefficient, and $\alpha$ is the diffusion exponent that characterizes the dynamical regime of the system. In particular, $\alpha = 1$ corresponds to normal (Brownian) diffusion, whereas $\alpha < 1$ indicates a subdiffusive regime associated with dynamical constraints or trapping effects in the condensate, and $\alpha > 1$ describes a superdiffusive behaviour. To obtain an accurate estimate of the MSD, the NVT trajectories of the condensates (see Section S4) and the same correlator technique used in other works for the calculation of the stress relaxation modulus were employed [17]. To obtain the diffusion coefficient $D$, we performed a least-squares fit when $\alpha = 1$. The monomeric MSD was obtained by tracking the position of each monomer over time using the same expression. Additionally, we estimate the diffusion time as the time at which at least 90% of the monomeric MSDs have $\alpha \geq 0.9$. We compute the time autocorrelation function of the end-to-end vector defined as:

$$\phi(t) = \langle \mathbf{R}(t) \cdot \mathbf{R}(0) \rangle , \tag{S12}$$

where $\mathbf{R}(t)$ is the instantaneous end-to-end vector of the protein at time $t$. In addition to the end-to-end definition, we also consider an alternative intrachain vector connecting one terminus to a residue located near the midpoint of the sequence (end-to-middle vector $\mathbf{R}_{1/2}(t)$). The corresponding autocorrelation function is computed analogously by replacing $\mathbf{R}(t)$ by $\mathbf{R}_{1/2}(t)$ , defined as $\mathbf{R}_{1/2}(t) = \mathbf{r}_0(t) - \mathbf{r}_{N/2}(t)$, or equivalently $\mathbf{R}_{1/2}(t) = \mathbf{r}_N(t) - \mathbf{r}_{N/2}(t)$, where $\mathbf{r}_0(t)$, $\mathbf{r}_N(t)$ and $\mathbf{r}_{N/2}(t)$ are the position of the first, last and middle amino acid, respectively. In the case of ProT$\alpha$ the end-to-middle relaxation functions results significantly symmetric to the centre so we averaged both functions (see Fig S1).

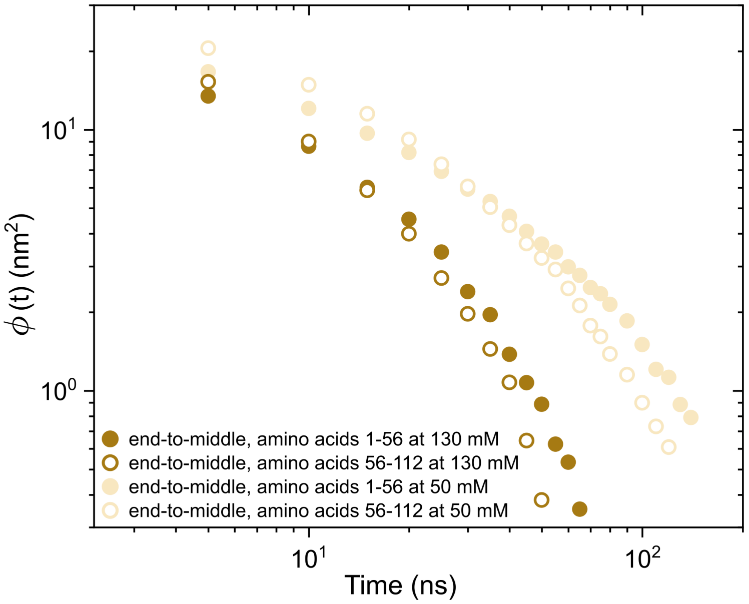


**FIG. S1:** End-to-middle relaxation functions for ProT$\alpha$ at 130 mM and 50 mM KCl. Solid symbols correspond to distances from amino acid 1 to 56, while open symbols correspond to distances from amino acid 56 to 112.

The autocorrelation function is computed from the NVT trajectories of the condensates under bulk conditions at 295 K, as described in Section S4. Specifically, $\mathbf{R}(t)$ and $\mathbf{R}_{1/2}(t)$ are evaluated for each protein molecule within the dense phase, and the correlation function is averaged over all chains and multiple time origins along the trajectory to improve statistical accuracy. To efficiently compute the relaxation function, we employ a multiple-$\tau$ correlator scheme [17, 21], which combines fine temporal resolution at short times with logarithmic sampling at long times. This approach enables accurate estimation of both fast local relaxation processes and the slow global reconfiguration dynamics. Considering that the relaxation of $\phi(t)$ can be expressed as a sum over normal modes, each associated with a characteristic relaxation time [22], we approximate the autocorrelation function using a truncated mode expansion involving the two slowest modes:

$$\phi(t) \approx A \exp\left(-\frac{t}{\tau_r}\right) + \frac{A}{4} \exp\left(-\frac{4t}{\tau_r}\right), \tag{S13}$$

where $\tau_r$ represents the longest (dominant) relaxation time, and $A$ is a prefactor related to the amplitude of the fluctuations. Thus, the reconfiguration time $\tau_r$ is obtained by fitting autocorrelation function to the expression above using a nonlinear least-squares procedure.

#### S8. CALCULATION AND ANALYSIS OF CONTACT MAPS

Intermolecular contact maps within protein condensates were calculated from the 3-5 $\mu$s NVT trajectories described in Section S4. Contacts were determined in all systems at 295 K and at the corresponding salt concentration. Typically, molecular contacts are identified based on a distance criterion, with the assumption that the relative frequency of contact map occurrences (rather than absolute frequency) remains generally unaffected by the selected cut-off distance used in calculations, provided the cut-off values are reasonable. However, to accurately capture the most relevant and common residue-residue contact pairs that facilitate the phase separation, it is highly recommended to account for the specific parameterisation of each amino acid in terms of excluded volume and minimum potential energy interaction distance. Hence, we adopted a sequence-dependent cut-off distance equivalent to $1.2\sigma_{ij}$, where $\sigma_{ij}$ represents the mean excluded volume of the respective i-th and j-th amino acid [23]. Given that the minimum of the potential used is approximately $\sqrt[6]{2}\sigma_{ij} \approx 1.122\sigma_{ij}$, we set the cut-off distance slightly beyond this point, at $1.2\sigma_{ij}$, to ensure significant binding. By implementing this sequence-dependent cut-off scheme for each amino acid pair interaction, we can effectively filter out adjacent contacts that may coincide with actual interacting amino acids along the sequence, thus enhancing our ability to accurately identify the amino acids that positively contribute to establishing condensates [23].

#### S9. CONTACT LIFETIMES ANALYSIS

To characterize the dynamical stability of intermolecular interactions within the condensates, we compute the distribution and characteristic timescale of inter-protein contact lifetimes from dedicated NVT trajectories at 295 K and the corresponding salt concentration, each comprising 500 ns of pre-equilibration followed by 30 ns of production sampled every 1 ps. Each contact, uniquely identified by a residue pair $(i, j)$ and monitored across consecutive snapshots, is defined using a transition-based (core-state) criterion [24]: two residues are considered to become newly bound once their separation first drops below $1.2\sigma_{ij}$, and the resulting contact is taken to persist, irrespective of brief excursions above this threshold, until the separation exceeds an outer distance of $1.8\sigma_{ij}$. This hysteresis between the formation and rupture thresholds, together with a one-frame tolerance for sub-resolution recrossings of the outer threshold, ensures that a recorded rupture corresponds to genuine dissociation rather than a thermal fluctuation about a single distance cutoff, which would otherwise systematically underestimate the lifetime and obscure differences in binding strength between systems. At each frame, contacts are accordingly classified as new, maintained, or broken; the lifetime of a completed contact is the number of consecutive frames for which it remained bound between formation and rupture, and contacts are grouped according to the interacting chain-length pair $(N_i, N_j)$ to allow comparison between systems (see main text for the resulting distributions).

The mean contact lifetime $\langle\tau\rangle$ for each chain-pair type is obtained as the ratio of the total time spent in the bound state to the number of completed rupture events. This estimator remains well defined even for the small fraction of contacts still bound at the end of the trajectory (right-censored events), whose observed bound time is retained in the numerator while contributing no rupture event to the denominator; across all systems this censored fraction is below 0.1%, indicating that $\langle\tau\rangle$ is a converged estimate rather than a lower bound set by the accessible simulation time. As an independent, censoring-free check of this timescale, we additionally compute the intermittent contact autocorrelation function $C(t) = \langle h(0)h(t)\rangle/\langle h\rangle$, where $h(t) = 1$ if a given residue pair is in the bound core state at time $t$ and 0 otherwise, averaged over all pairs of a given chain-length type and over time origins along the trajectory. Because $C(t)$ does not require a contact to have already ended, it carries no censoring bias by construction, at the cost of retaining a memory of intermittent rebinding that the completed-lifetime estimator $\langle\tau\rangle$ does not; Fig. S2 reports $C(t)$ together with the corresponding $\langle\tau\rangle$ values for the H1+ProT$\alpha$ and Protamine+ProT$\alpha$ systems at 50 and 160 mM salt.

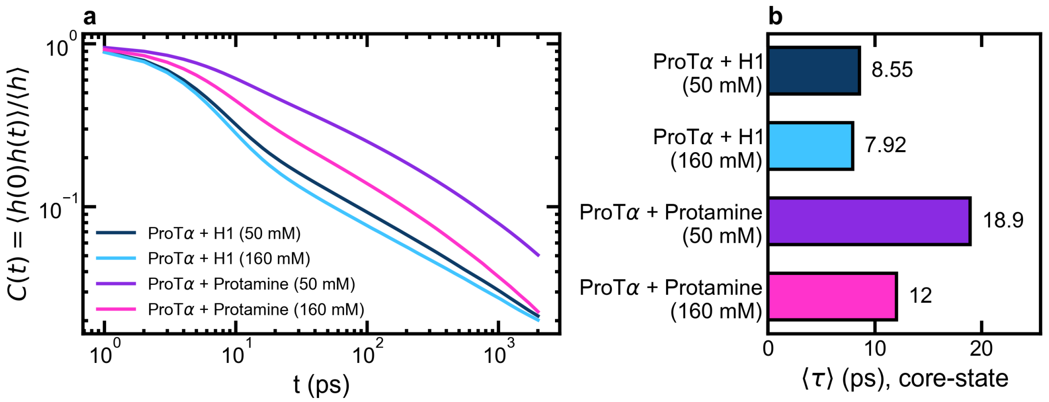


**FIG. S2:** (a) Intermittent contact autocorrelation function $C(t)$ for ProT$\alpha$-H1 (dark navy, 50 mM; light blue, 160 mM) and ProT$\alpha$-protamine (purple, 50 mM; magenta, 160 mM) at 295 K. (b) Corresponding mean contact lifetimes $\langle\tau\rangle$, obtained as the ratio of total bound time to the number of completed rupture events for each system.

#### S10. CONTACT MAPS OF PROT$\alpha$ IN THE COMPLEX COACERVATES

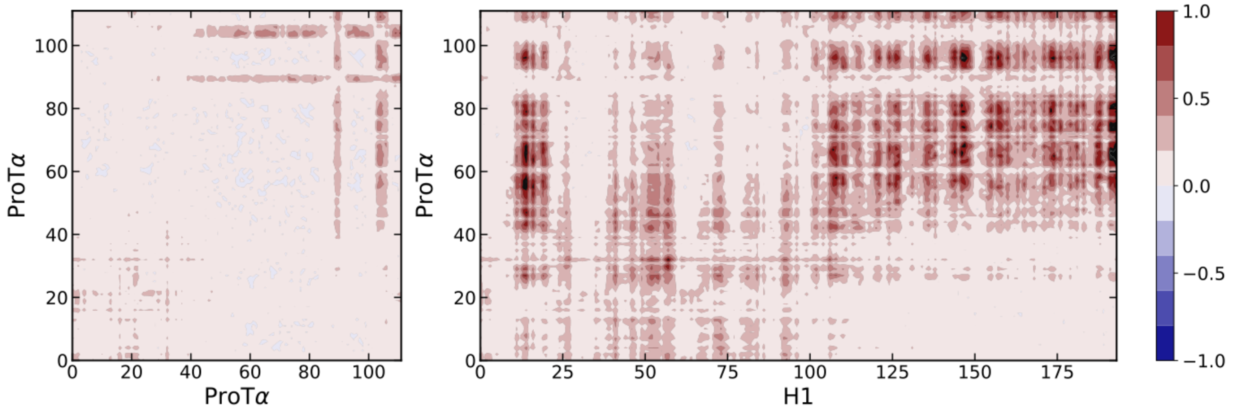


**FIG. S3:** Intermolecular contact frequency difference (in number of contacts per residue) between ProT$\alpha$-ProT$\alpha$ (left) and H1-ProT$\alpha$ (right) between the system at 50 and 160 mM of KCl.

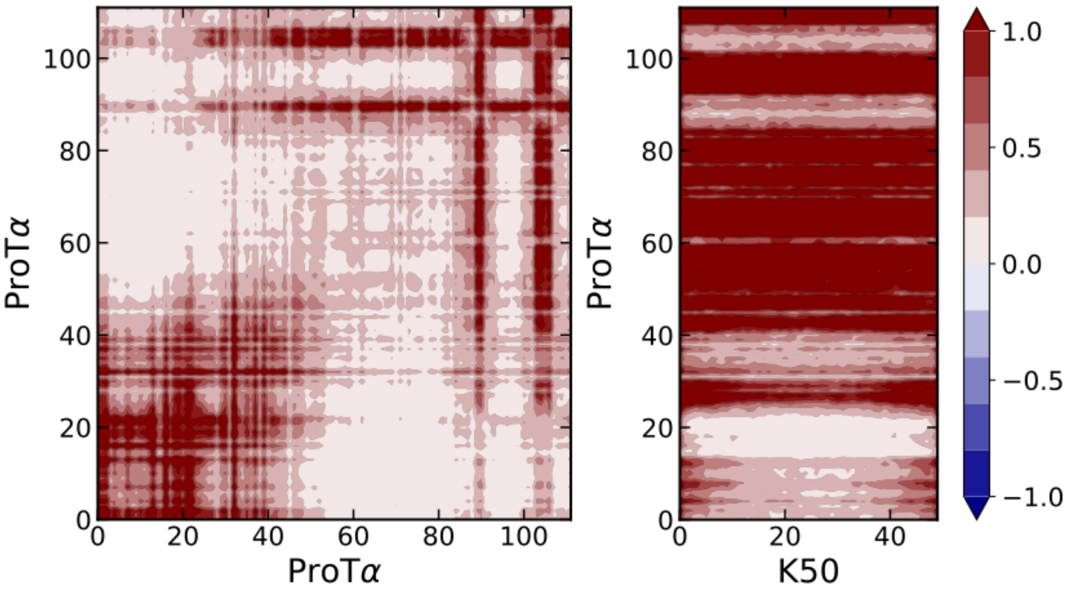


**FIG. S4:** Intermolecular contact frequency difference (in number of contacts per residue) between ProT$\alpha$-ProT$\alpha$ (left) and K50-ProT$\alpha$ (right) between the system at 50 and 210 mM of KCl.

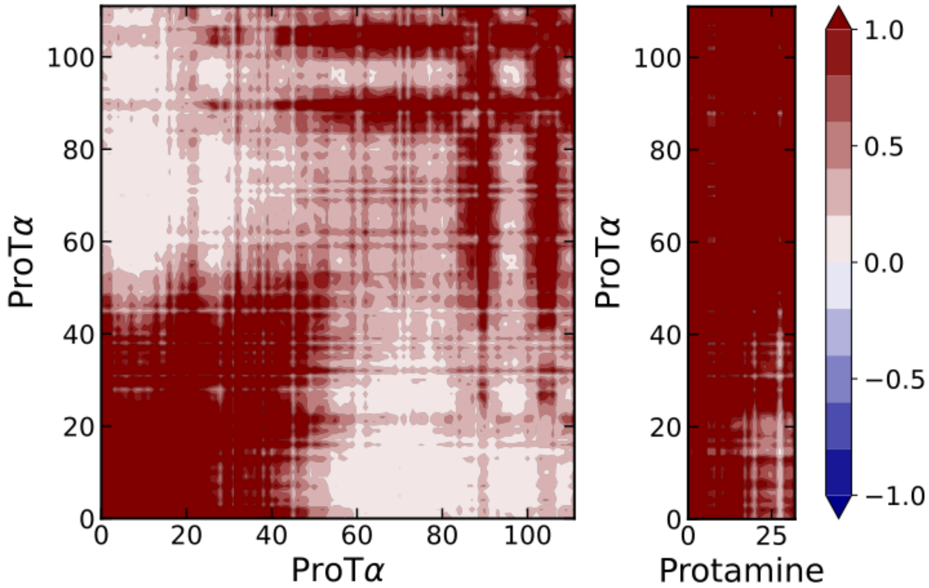


**FIG. S5:** Intermolecular contact frequency difference (in number of contacts per residue) between ProT$\alpha$-ProT$\alpha$ (left) and Protamine-ProT$\alpha$ (right) between the system at 50 and 250 mM of KCl.

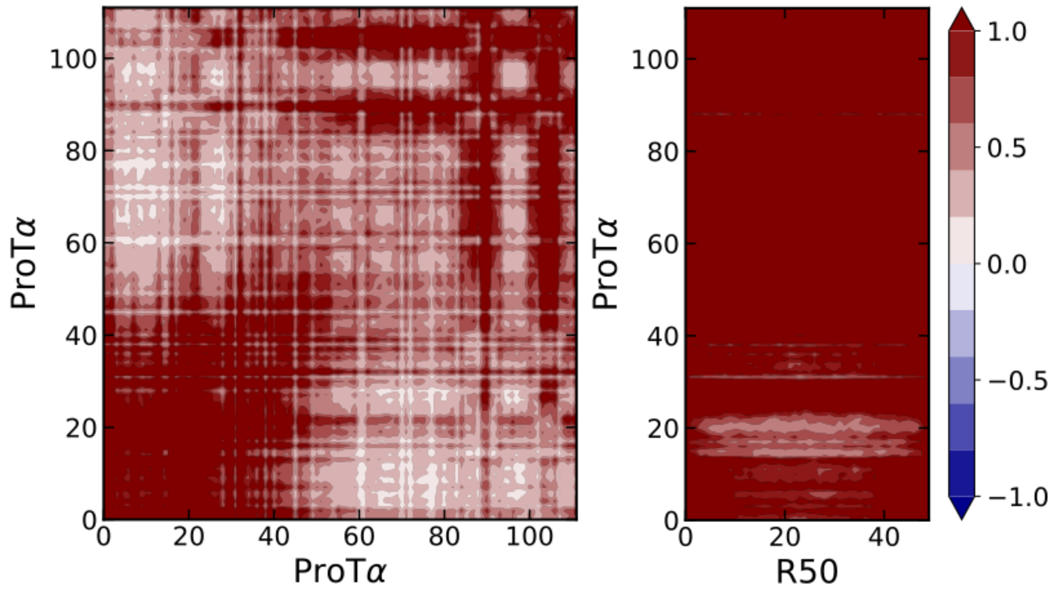


**FIG. S6:** Intermolecular contact frequency difference (in number of contacts per residue) between ProT$\alpha$-ProT$\alpha$ (left) and R50-ProT$\alpha$ (right) between the system at 110 and 400 mM of KCl.

#### S11. ANALYSIS OF THE NETWORK CONNECTIVITY BY A PRIMITIVE PATH ANALYSIS

We analyse the network connectivity of the complex coacervates using the NVT trajectories (see Section S4) and the primitive path analysis (PPA) method [25], modified in Tejedor *et al.* [26]. The PPA algorithm minimizes the contour length of all chains in the condensate while keeping the terminal residues of the monomers fixed and, in this way, preserving the network topology by preventing chains from crossing each other. The energy minimization is performed with a tolerance of $10^{-7}$ for both the energy and the force, allowing up to $2 \times 10^5$ iterations. During the minimization, several structural topological observables are computed considering chains with lengths within the specified range. In particular, the mean squared end-to-end distance $\langle \mathbf{R}^2 \rangle$ and the primitive path contour length $L_{pp}$ are extracted. The tube diameter $a$ is defined through the relation $\langle \mathbf{R}^2 \rangle = a \cdot L_{pp}$ [27]. The average number of entanglements per chain is then estimated as $Z = L_{pp}/a$.

#### S12. EXTENDED DISCUSSION OF RECONFIGURATION AND DIFFUSION TIME DEFINITIONS

The extracted reconfiguration time depends on the segment of the protein sequence used to define the distance in the autocorrelation function. In addition to the full end-to-end vector (Section S7), we compute end-to-middle autocorrelation functions by averaging the correlation from each terminus to the central residue (Fig. S1). As expected, end-to-middle correlations decay systematically faster than end-to-end correlations. Although both remain within the same order of magnitude, the reconfiguration times obtained from the end-to-middle correlations are ∼25% and ∼10% shorter than the end-to-end values at 50 and 130 mM, respectively, reflecting the intrinsic length-scale dependence of $\tau_r$. This indicates a potential nuance in experimentally obtained reconfiguration times, particularly in FRET measurements, which typically probe labelling positions that are not strictly at the terminal segments of the sequence [28].

The translational diffusion time $\tau_d$ is often determined from the mean squared displacement (MSD) of the protein centre of mass, $\mathrm{MSD}_{CM}(t) = \langle(\mathbf{r}_{CM}(t) - \mathbf{r}_{CM}(0))^2\rangle$. Given the difficulty of reliably capturing this parameter from the centre-of-mass MSD alone within accessible simulation times, we instead define $\tau_d$ as the time at which at least 90% of the monomeric MSDs have a local diffusive exponent $\alpha \geq 0.9$, marking the onset of the regime in which $\mathrm{MSD}(t > \tau_d) \propto t$. This corresponds to the longest relaxation time of the molecule [22, 29] and is associated with an average displacement on the order of the molecular size.

The stress relaxation modulus $G(t)$ (Section S6) exhibits two distinct regimes: a short-time regime dominated by fluctuations in short-range interactions and internal conformational rearrangements, and a long-time regime governed by the relaxation of intermolecular forces, large-scale conformational changes, and translational diffusion of proteins across the condensate [23, 30]. At very low salt concentrations, moderate discrepancies between simulated and experimental viscosities may arise from limited sampling, since the associated long relaxation times require simulation timescales exceeding $\sim 5\ \mu$s to observe the full decay of $G(t)$.

#### S13. EXTENDED RESIDUE-LEVEL CONTACT DYNAMICS

The temporal evolution of intermolecular contacts reveals rapid rearrangement of electrostatic pairings within the main binding hotspots identified in the contact maps (Section S10). In the ProT$\alpha$-H1 mixture, residues E63-E65 of ProT$\alpha$ initially interact with K161 of H1 and subsequently switch to K158 within $\sim$2 ps, indicating local charge-mediated reorganization within the primary hotspot spanning residues 55-85. A second hotspot is formed by acidic residues around positions 90-100 of ProT$\alpha$. In the ProT$\alpha$-protamine system, residues D95-D96 establish contacts with R15-R17, consistent with the contact maps of Section S10, evolving from an initial D96-R16 interaction toward D95-R16 and D96-R17 pairings within $\sim$2 ps. These observations illustrate the rapid and continuous reorganization of electrostatic intermolecular contacts at intermediate timescales within the condensate, underlying the short average contact lifetimes reported in Section S9.

For a typical homopolymer, the monomeric MSD at short times is symmetric with respect to the central monomer (see Fig. 2B in Ref. [31]). In contrast, the sequence-specific charge patterning of ProT$\alpha$ strongly breaks this symmetry: residues in the 60-80 segment, and to a lesser extent around 90-100, move significantly slower than the rest of the chain at short times, consistent with their involvement in persistent electrostatic contacts, and this profile reproduces the key features reported in atomistic simulations [24].

#### S14. EXTENDED POLYMER-PHYSICS FRAMEWORK FOR CONDENSATE DYNAMICS

Chain dynamics at the local scale in these condensates are governed by a large number of low-energy, short-lived residue-residue interactions, whose lifetimes are several orders of magnitude shorter than the conformational reconfiguration time and translational diffusion time. Such regimes can be described using two complementary polymer frameworks [24]: the sticky Rouse model [32, 33] and the Rouse model with entanglements based on reptation theory [22, 34, 35]. At higher polymer densities, topological constraints give rise to entanglement effects beyond the Rouse regime, described within reptation theory as motion confined to a tube imposed by surrounding chains [34, 36–38]; extensions incorporating contour length fluctuations enable a more realistic description of dense systems [22, 39]. Given the chain lengths and densities of the proteins studied here, the dynamics are expected to lie in an intermediate regime between the Rouse and entangled limits [24, 40–42].

For reference, in the coarse-grained Kremer-Grest model [43], a benchmark framework for MD simulations of polymer melts, chains typically require $N \gtrsim 140$ beads to reach the entangled regime ($Z \sim 1.7$) at a monomer density of 0.85 [27, 41]. The monomer density in our condensates spans 0.27 to 1.38 (assuming an average bead size $\sigma = 0.65$ nm and an average residue mass $m = 120$ g/mol), placing the systems near the crossover between the Rouse and weakly entangled regimes [40, 41], consistent with the entanglement numbers $Z \approx 1.2$-2.5 obtained from primitive path analysis (Section S11).

The remaining quantitative deviations between the simulated viscosity, diffusion coefficient and reconfiguration time and the power laws expected from Rouse/reptation theory (main text) can be attributed to the proximity of the systems to the crossover between the two regimes, where topological constraints introduce corrections proportional to $\langle R^2 \rangle / a^2$. These results support a physical picture in which condensate dynamics are predominantly Rouse-like, with moderate corrections arising from transient entanglements, leading to a transition from low- to high-viscosity states as a function of salt concentration.

#### S15. EXTENDED RESULTS ON CLUSTER FORMATION AND NUCLEATION

For each salt concentration and protein density, the nucleation free energy as a function of cluster size, $\Delta G(N) = -\ln(n_N / N_{\text{chains}})$ (in units of $k_B T$, referenced to the monomer, $N = 1$), is obtained from the population $n_N$ of clusters of size $N$ observed in the pre-nucleation portion of each trajectory—the initial fraction of frames preceding sustained cluster growth—normalized by the average number of chains per frame. This is the same cluster-size-distribution approach used to resolve nucleation free-energy landscapes in electrolyte solutions [44]. Cluster sizes for which the normalized population falls below $5 \times 10^{-4}$ of the monomer population are considered too poorly sampled for a reliable estimate and are excluded (blank cells in Fig. 6c of the main text).

At low protein density, cluster formation strongly depends on salt concentration. At 60 mM, chains progressively associate into clusters that grow at a slow but steady pace, reaching ~150 chains by the end of the 2 $\mu$s trajectory (Fig. 6a of the main text), reflecting reduced electrostatic screening, which favours persistent protein-protein association even at low concentration. At 150 and 200 mM, no stable clusters are observed at this density, and chains remain predominantly free in solution. At high protein density, close to the critical density, the three salt conditions lead to markedly different outcomes: at 60 mM, chains coalesce into a single, large cluster comprising nearly all 200 chains of the system, indicative of a highly stable, percolated condensate; at 150 mM, clusters remain smaller and more heterogeneous, fluctuating between approximately 50 and 100 chains; at 200 mM, large clusters do not form even at this higher density, indicating that electrostatic screening is sufficient to substantially destabilize ProT$\alpha$-H1 association across the entire density range explored.

The cluster size distribution in the pre-nucleation state reveals that, under all conditions studied, a dimer is more likely to be found than a trimer or a higher-order complex, consistent with the greater stability of the PH complex reported from the PMF calculations (Section S5). At 60 mM this maximum is not clearly resolved because the system nucleates rapidly, hindering sampling of the pre-nucleation state; at 150 and 200 mM, where nucleation is slower, the maximum around the dimer is clearly observed.

The nucleation free-energy barrier for a cluster of size $N$ increases with salt concentration, and for high values of $N$, less dense systems display higher barriers than denser ones, consistent with the lower probability of finding chains in close proximity at low density. For reference clusters of $N_{ref} = 5$ chains, the nucleation energy at 60 and 150 mM is approximately constant and independent of system density, whereas at 200 mM the barriers are systematically higher, in good agreement with the stronger electrostatic screening disfavouring chain association. For $N_{ref} = 10$ chains, density has a much more pronounced influence: at 60 mM the barriers are relatively low across the entire density range explored; at 150 mM, overcoming the barrier is considerably more difficult; and at 200 mM the barriers are the highest of the three salt concentrations and grow as density decreases, while at the two lowest densities sampling is insufficient to reliably determine them.

[31] A. Castro, J. Luengo-Marquez, A. R. Tejedor, R. Collepardo-Guevara, M. Papp, P. Arosio, A. Ocana, I. Sanchez-Burgos, and J. Rene Espinosa, "Interplay between local diffusion, concentration, and inter-protein alignment promotes cross-$\beta$-sheet transitions at condensate interfaces," *bioRxiv*, 2026.

[32] M. Rubinstein and A. N. Semenov, "Dynamics of entangled solutions of associating polymers,"

*Macromolecules*, vol. 34, no. 4, pp. 1058–1068, 2001.
